# Alien leaf beetles of European Russia: native ranges, invasion history, biology and vectors of dispersal

**DOI:** 10.1101/252510

**Authors:** Andrzej O. Bieńkowski, Marina J. Orlova-Bienkowskaja

## Abstract

Invasions of leaf beetles are of great ecological and economic significance, but poorly studied. The rate of these invasions in Europe is dramatically increasing. Some established species spread quickly occupying almost the whole continent within several decades. We present the first inventory of alien leaf beetles of European Russia. For each species the map of distribution is provided and the history of invasion in the world is discussed. Two species native to Mediterranean Region: *Chrysolina americana* (pest of *Rosmarinus* and *Lavandula)* and *Leptomona erythrocephala* feeding on *Lotus corniculatus* are recorded in European Russia for the first time. A polyphagous pest of floriculture *Luperomorpha xanthodera* native to China and Korea and pest of soybeans *Medythia nigrobilineata* native to East Asia were recorded in 2016. A pest of tobacco *Epitrix hirtipennis* native to North America was recorded in 2013. A pest of corn *Diabrotica virgifera* was intercepted at the border of Russia in 2011, but is not established. Three alien species were recorded in the 20th century: *Zygogramma suturalis* introduced from North America for control of *Ambrosia, Phyllotreta reitteri* native to Afghanistan and Tajikistan and feeding on *Lepidium latifolium*, and the Colorado potato beetle *Leptinotarsa decemlineata*. The Black Sea region is more prone to leaf beetle invasions than other regions of European Russia. Leaf beetles usually occur only on alien or cultivated plants. Some species feed on native plants in native communities. So it is difficult to distinguish species established before the 20th century from native ones.

## Introduction

There are major gaps in knowledges on regional alien floras and faunas and data availability varies among regions (Pyšek et al. 2017). In particular, alien insects of European Russia are poorly studied. The most of information is published in local sources and stays unknown to the wide audience, because is not included to international databases. We present the first inventory of alien leaf beetles of this territory.

Leaf beetles (Chrysomelidae except Bruchinae) is a large group of phytophagous beetles (more than 36 000 species in the world, about 600 in European Russia) (Jolivet 1995; Bieńkowski 2011). Negative economic impact of invasions of these insects is significant, since some of them, e.g. *Leptinotarsa decemlineata, Diabrotica virgifera* and *Epitrix papa*, are major pests of various crops (EPPO 2017). In spite of it, invasions of leaf beetles are poorly studied (Beenen 2006). Before 2010 only 11 species alien to Europe were recorded (Beenen and Roques 2010), and the information about invasions of leaf beetles to European Russia was restricted to *Leptinotarsa decemlineata* and *Zygogramma suturalis* (Ivanchik and Izhevsky 1981; Kovalev and Medvedev 1983; Maslyakov and Izhevsky 2011).

Four alien species were first recorded in European Russia in 2011–2016: *Diabrotica virgifera* (VNIIKR 2012), *Epitrix hirtipennis* (Orlova-Bienkowskaja 2014), *Luperomorpha xanthodera* (Bieńkowski and Orlova-Bienkowskaja 2018a) and *Medythia nigrobilineata* (Bieńkowski and Orlova-Bienkowskaja 2018b). Analysis of ecology and dynamics of ranges of some other species has revealed that they could represent archeoinvaders, i.e. alien species established in European Russia before the 20th century (Orlova-Bienkowskaja 2013a, b). Here we present the first records of two alien species new to European Russia: *Leptomona erythrocephala* and *Chrysolina americana* and the review of all other leaf-beetles alien to this region. For each species a database of records and a map of geographical distribution are compiled and the following information is provided: data on host plants and other features of biology, description of native range, invasion history, possible vectors of dispersal and economic impact of establishment. This article is a part of the project “Alien beetles of European Russia”.

## Materials and methods

Distribution of alien species was studied on the base of own collection of the authors in European Russia in 1988–2017, examination of specimens from collections of ZIN (Zoological Institute of Russian Academy of Sciences, St. Petersburg, Russia), ZMMU (Zoological Museum of Moscow State University, Moscow, Russia), FNMS (Senckenberg Naturmuseum, Frankfurt am Main, Germany), HNHM (Hungarian Natural History Museum, Budapest, Hungary), MTD (Museum für Tierkunde, Dresden, Germany), NHMW (Naturhistorisches Museum, Wien, Austria), NMP (National Museum in Prague, Prague, Czech Republic), VNIIKR (Russian Plant Quarantine Center, Bykovo, Russia), BSU (Belgorod State University, Belgorod, Russia), OSU (Orel State University, Orel, Russia), BNR (Belogorie Nature Reserve, Borisovka, Russia), and 17 private collections in Russia: AB (A.O. Bieńkowski, Zelenograd), AK (A.G. Koval, St. Petersburg), AP (A.I. Prikhodko, Zelenograd), AR (A.B. Ruchin, Saransk), AU (A.S. Ukrainsky, Moscow), EI (E.V. Iljina, Makhachkala), GK (G.A. Korostov, Elista), MD (M.M. Dolgin, Syktyvkar), NN (N.E. Nikolaeva, Tver), NO (N.V. Okhrimenko, Krasnodar), PP (P.N. Petrov, Moscow), PR (P.V. Romantsov, St. Petersburg), RI (R.N. Ishin, Tambov), SM (S.A. Mosyakin, Simferopol), TM (T.A. Mogilevich, Zelenograd), VF (V.I. Filippov, Sochi), YK (Y.N. Kovalenko, Moscow). In addition data from Global Biodiversity Information Facility and from 82 literature sources were used. We use criteria of alien status and terminology used in invasion biology defined by Richardson et al. (2000) and Blackburn et al. (2011).

Here we adhere to the system adopted in the Catalogue of Palaearctic Coleoptera (Kippenberg 2010; Beenen 2010; Döberl 2010). We study invasions only of leaf-beetles in the narrow sense of the word, i.e. all subfamilies of Chrysomelidae except seed beetles (Bruchinae), because seed beetles are very different from other Chrysomelidae in biology and ecology and have different vectors and trends of invasions (Beenen and Roques 2010). Species below are presented in reverse chronological order of their first records in European Russia.

### 2017 – *Leptomona erythrocephala* (Olivier, 1790) (Galerucinae)

#### Native range

*Leptomona erythrocephala* is native to mainland Spain, Mallorca, Portugal, south France, Sicily, Algeria and Morocco (Fig. 1) (Reitter 1886; Biondi et al. 1995; Beenen 2010; Aberlenc 2010). Junior synonym *Monolepta verticalis* Reitter, 1886 was described from Portugal (Reitter 1886). *Leptomona erythrocephala* was also recorded from northern Italy, but this record is supposed to be questionable (Biondi et al. 1995). Database on localities of *L. erythrocephala* is provided in electronic supplementary material 1.

**Fig. 1.**
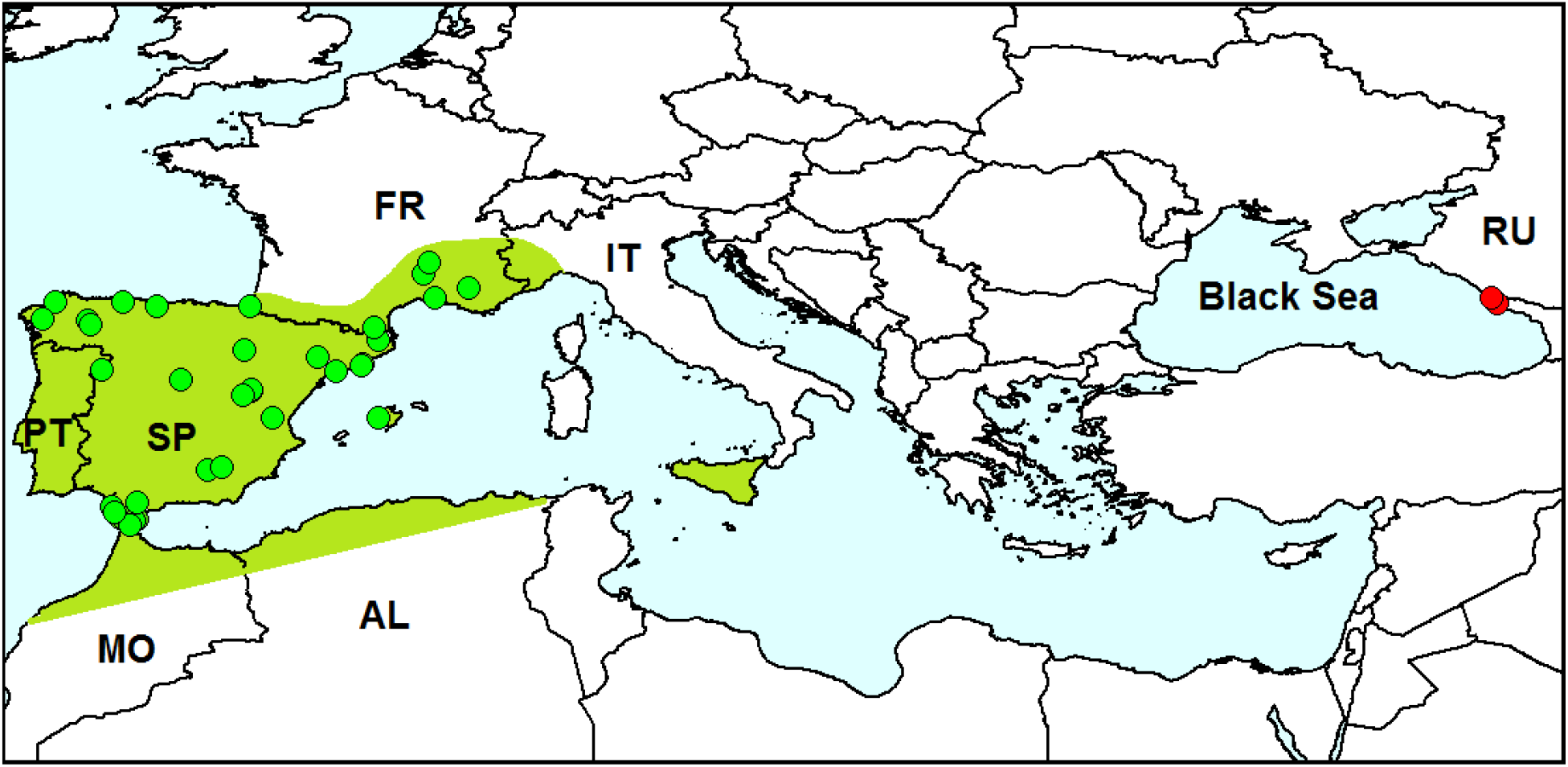
Distribution of *Leptomona erythrocephala. Red dots* – localities of specimens outside species native range (original data), *green dots* on the *green background* – localities of specimens in the native species range. AL – Algeria, FR – France, IT – Italy, MO – Morocco, PT – Portugal, RU – Russia, SP – Spain. Sources of information about localities of the species in its native range: Heyden and Allard (1870), Reitter (1886), Champion (1902), Daccordi and Petitpierre (1977), Petitpierre (1981, 1997, 2005, 2007), Aberlenc (1987, 2010), Biondi et al. (1995), García-Ocejo and Gurrea (1995), Baselga and Novoa (2003, 2004, 2006), Staines and Whittington (2003), Schmidt et al. (2005), Petitpierre et al. (2011, 2017), Petitpierre and Daccordi (2013), GBIF (2017a).

#### Invasion history

This species was not recorded outside its native range before.

#### Records in European Russia

In 2017 we found *L. erythrocephala* in the north-west Caucasus, near the Black Sea coast. The first specimen was found on June 3 on the bank of the pond in Imeretian resort by sweep-netting (43°24’N, 39°58’E). Then on June 6 about 50 specimens were found on the ruderal plant *Lotus corniculatus* in Razdolnoe, in the floodplain of Bzugu River (43°36’N, 39°46’E).

#### Taxonomy and identification

The genus *Leptomona* Bechyné, 1958 was distinguished by J. Bechyné from the large genus *Monolepta* Chevrolat, 1836 and includes four species: *L. erythrocephala* distributed in Mediterranean region, *L. russica* (Gmelin, 1790) distributed in the steppes of Eastern Europe and western Asia, *L. fulvicollis* (Jacoby, 1885) distributed in Japan and *L. subseriata* (Weise, 1887) distributed in East Siberia and the Far East. Only one species – *L. russica* has been recorded in European Russia (Bieńkowski and Orlova-Bienkowskaja 2013).

*Leptomona erythrocephala* differs from other species of the genus by the shape of aedeagus (Fig. 2) and by the following characters: punctation of elytra entirely confused; pronotum covered with punctures which slightly smaller than those on elytra; head, pronotum, prosternum, coxae, femora, tibiae and antennomeres 1–3 reddish, mesosternum brown, elytra blue, labrum, tarsi, metasternum, abdominal tergites and sternites black; hind wings reduced. Identification of specimens collected in Sochi (see electronic supplementary material 2) was confirmed by their comparison with specimens of *L. erythrocephala* from ZIN collected in Spain.

**Fig. 2.**
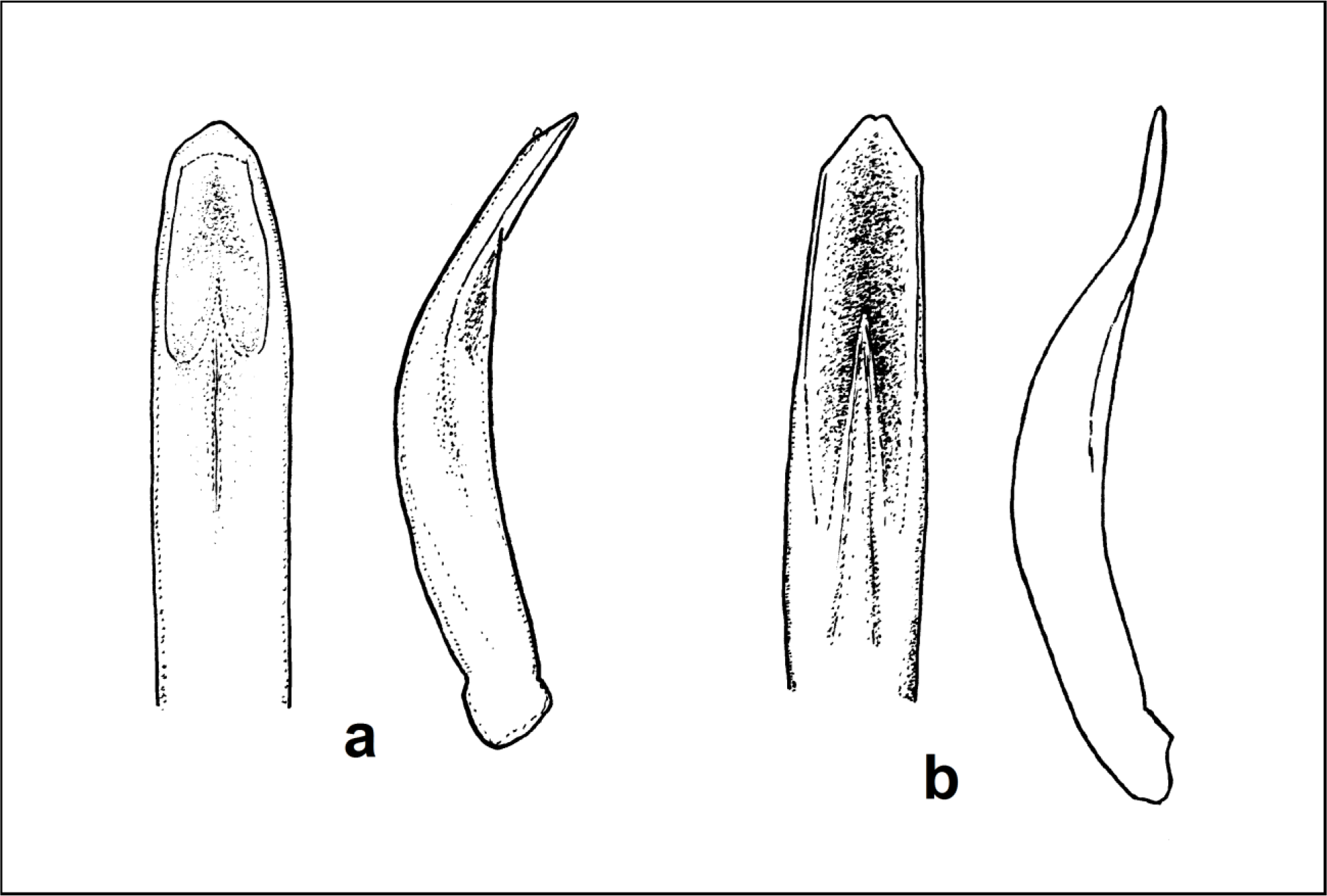
Aedeagus of *Leptomona erythrocephala* from Sochi (**a**) and *L. russica* (**b**).

#### Biology

Host plant of this species was unknown till now. But *L. russica* and representatives of the closely related genus *Monolepta* feed mainly on Fabaceae (Medvedev and Roginskaya 1988). Our observations in nature have shown that *L. erythrocephala* feeds on *Lotus corniculatus* (Fabaceae), which is a common ruderal plant in the city. Feeding on this plant was confirmed by our observations in cage. Ten specimens were placed in cage with the following Fabaceae plants collected in the same biotope: *Trifolium repens, Trifolium aureum, Gleditsia triacanthos* and *Lotus corniculatus*. The beetles fed only on *L. corniculatus:* gnawed margins of leaves and petals.

#### Invasion status

Obviously, *Leptomona erythrocephala* is established in the region. First, many specimens were collected in the wild in two localities. Second, it feeds on the native plant.

#### Vector of dispersal

Sochi is more than 2000 km far from the native range of the species. So the natural spread is impossible. The most probable dispersal vector is an unintentional introduction with planting material or soil. The distance between two localities where *L. erythrocephala* was found in the Caucasus is 28 km. Since *L. erythrocephala* is a flightless beetle and its natural dispersal ability is restricted, it seems that dispersal of *L. erythrocephala* in the region is connected with an unintentional introduction by man.

#### Economic impact

*Leptomona erythrocephala* is not regarded as a pest in its native range. But special attention should be payed to trophic specialization of this species in the Caucasus, especially if it becomes abundant. It should be taken into account that the representatives of the genera *Leptomona* and *Monolepta* feed on Fabaceae, and one of them, *Monolepta quadriguttata* (Motschulsky, 1860), is a serious pest of soybean (Medvedev and Roginskaya 1988; Moseiko 2010).

### 2016 – *Luperomorpha xanthodera* (Fairmaire, 1888) (Alticinae)

#### Native range

Rose flea beetle is native to China and Korean peninsula (Döberl 2010).

#### Invasion history

(Fig. 3). This pest was firstly found in Europe in 2003 on British Islands (Johnson and Booth 2004). Then it quickly spread to Italy (Conti and Raspi 2007), France (Doguet 2008), Germany, Switzerland (Döberl and Sprick 2009), The Netherlands (Beenen et al. 2009), Hungary (Bodor 2011), Austria (Geiser and Bernhard 2012), Poland (Kozłowski and Legutowska 2014), Belgium (Fagot and Libert 2016), Spain (Viñolas et al. 2016) and European Russia (Bieńkowski and Orlova-Bienkowskaja 2018a).

**Fig. 3.**
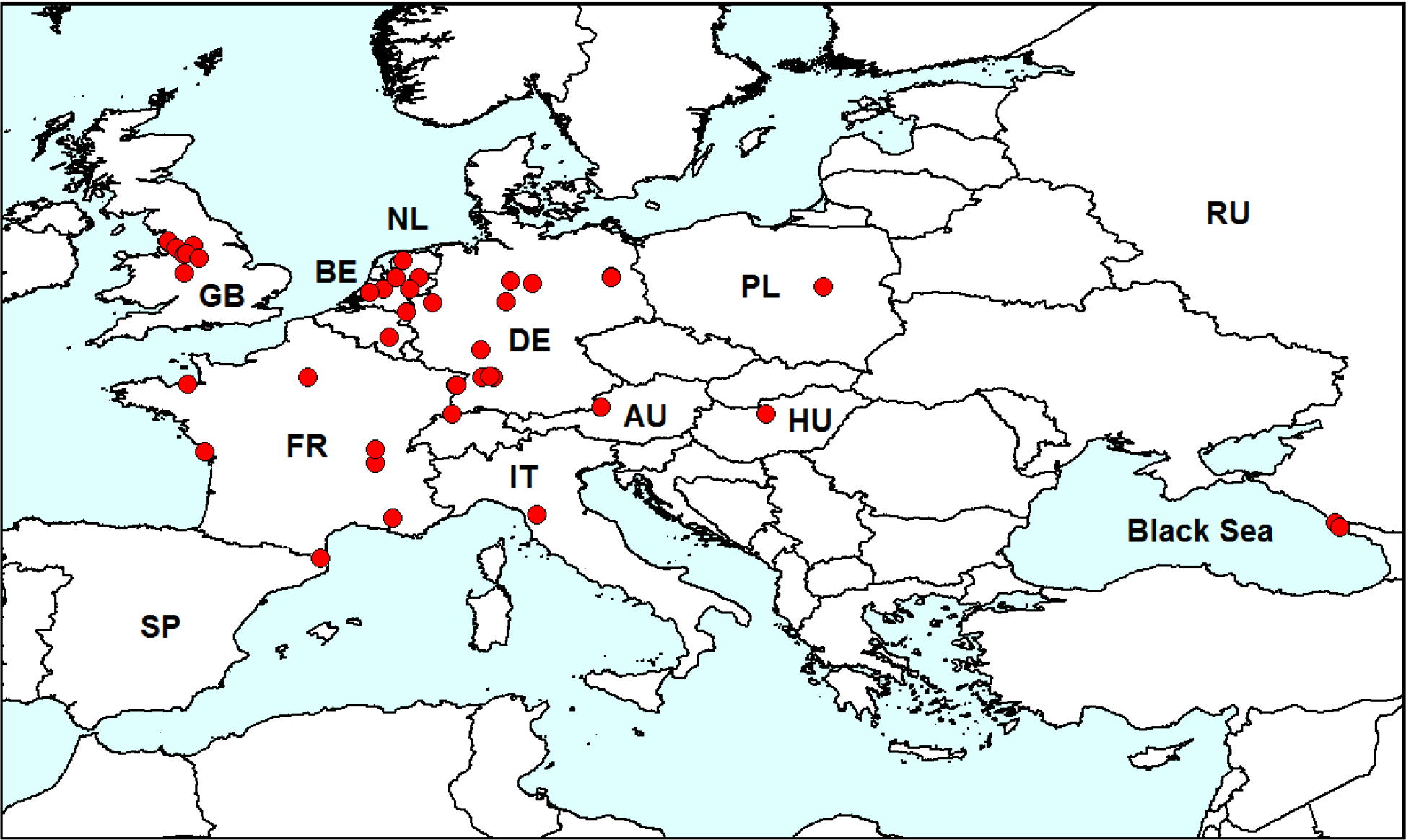
Distribution of *Luperomorpha xanthodera* in Europe. *Red dots* – localities of *L. xanthodera* in Europe. AU – Austria, BE – Belgium, DE – Germany, FR – France, GB – Great Britain, HU – Hungary, IT – Italy, NL – Netherlands, PL – Poland, RU – Russia, SP – Spain. Sources of information: Johnsonl and Booth (2004), Doguet (2008), Conti and Raspi (2007), Döberl and Sprick (2009), Beenen et al. (2009), Vincent and Doguet (2011), Bodor (2011), Geiser and Bernhard (2012), Kozłowski and Legutowska (2014), Fagot and Libert (2016), Viñolas et al. (2016), Callot (2017), Heinig and Schöller (2017), Bieńkowski and Orlova-Bienkowskaja (2018a) and original data (see below). Database on localities of *L. xanthodera* is provided in electronic supplementary material 3.

#### Records in European Russia

In 2016 *L. xanthodera* was firstly recorded in European Russia (Bieńkowski and Orlova-Bienkowskaja 2018a). On 22.05.2016 we found one female in Adler district of Sochi in the wastelands and roadside at south-west border of the international airport Sochi (Adler) (43°26’N, 39°55’E). On 23.05.2016 numerous beetles were observed in the Central district of Sochi (43°35’N, 39°45’E) on rose flowers. On 8.06.2017 two males and one female were collected with a sweep net in Adler district of Sochi, in Olympic Village on grasses on the bank of the pond (43°25’N, E 39°56’E).

#### Biology

*Luperomorpha xanthodera* is a polyphagous species. Adults feed on flowers of many plants (23 genera from 19 families), larvae develop on roots of these plants (Del Bene and Conti 2009). In Sochi adults of *L. xanthodera* feed flowers of roses and citrus plants (Bieńkowski and Orlova-Bienkowskaja 2018a).

#### Vector of dispersal

*Luperomorpha xanthodera* was probably unintentionally introduced as larvae on roots of imported seedlings, or as adults transported as cargo stowaways in airplanes. Both vectors are quite possible in the region of Sochi. First, this city is close to the international airport. Second, mass planting with imported planting material was carried out there during the landscaping of the city in preparation for the Olympic Games, which took place in 2014. Many other invasive pests have been introduced and established in the city in this period (Bieńkowski and Orlova-Bienkowskaja 2018a).

#### Invasion status

*Luperomorpha xanthodera* is established in the region. Finds of numerous specimens in two subsequent years in different localities indicate, that a self-sustaining population exists in the wild, and the species is dispersing in the region.

#### Economic impact

Establishment of *L. xanthodera* in the south of European Russia could cause negative economic consequences, since it is a pest of ornamental flowers.

### 2016 – *Medythia nigrobilineata* (Motschulsky, 1861) (Galerucinae)

#### Native range

Two-striped leaf beetle *Medythia nigrobilineata* (= *Paraluperodes suturalis nigrobilineatus)* is native to North China, Japan, Nepal, Pakistan, South Korea, East Siberia and the Russian Far East (Fig. 4) (Medvedev 1992; Beenen 2010; Warchałowski 2010; Toepfer et al. 2014).

#### Invasion history

In 2016 this species was firstly found outside its native range - in the south of European Russia (Bieńkowski and Orlova-Bienkowskaja 2018b).

#### Record in European Russia

A single female specimen of *Medythia nigrobilineata* was collected by sweep-netting on grass (43°25’N, 39°59’E) on May, 19, 2016 in Krasnodar Krai, city of Sochi, near Adler, Imereti lowland, on wasteland with grasses (Bieńkowski and Orlova-Bienkowskaja 2018b).

#### Biology

Biology of *M. nigrobilineata* in its native range was described by Ogloblin (1936), Koyama (1940), Moseiko (2010) and Toepfer et al. (2014). *Medythia nigrobilineata* develops only on soybeans. Adults feed on leaves and often damage immature pods. Besides that, beetles can feed on leaves of rice and sugarcane. Adults hibernate among fallen leaves and in the soil. In spring they begin to feed on soybean seedlings and damage leaves. Females lay eggs in the soil. Larvae feed on root nodules and pupate in the soil.

#### Vector of dispersal

The specimen was collected 4 km from the international airport Sochi. We suspect that *M. nigrobilineata* could be unintentionally introduced from Asia through this airport.

**Fig. 4.**
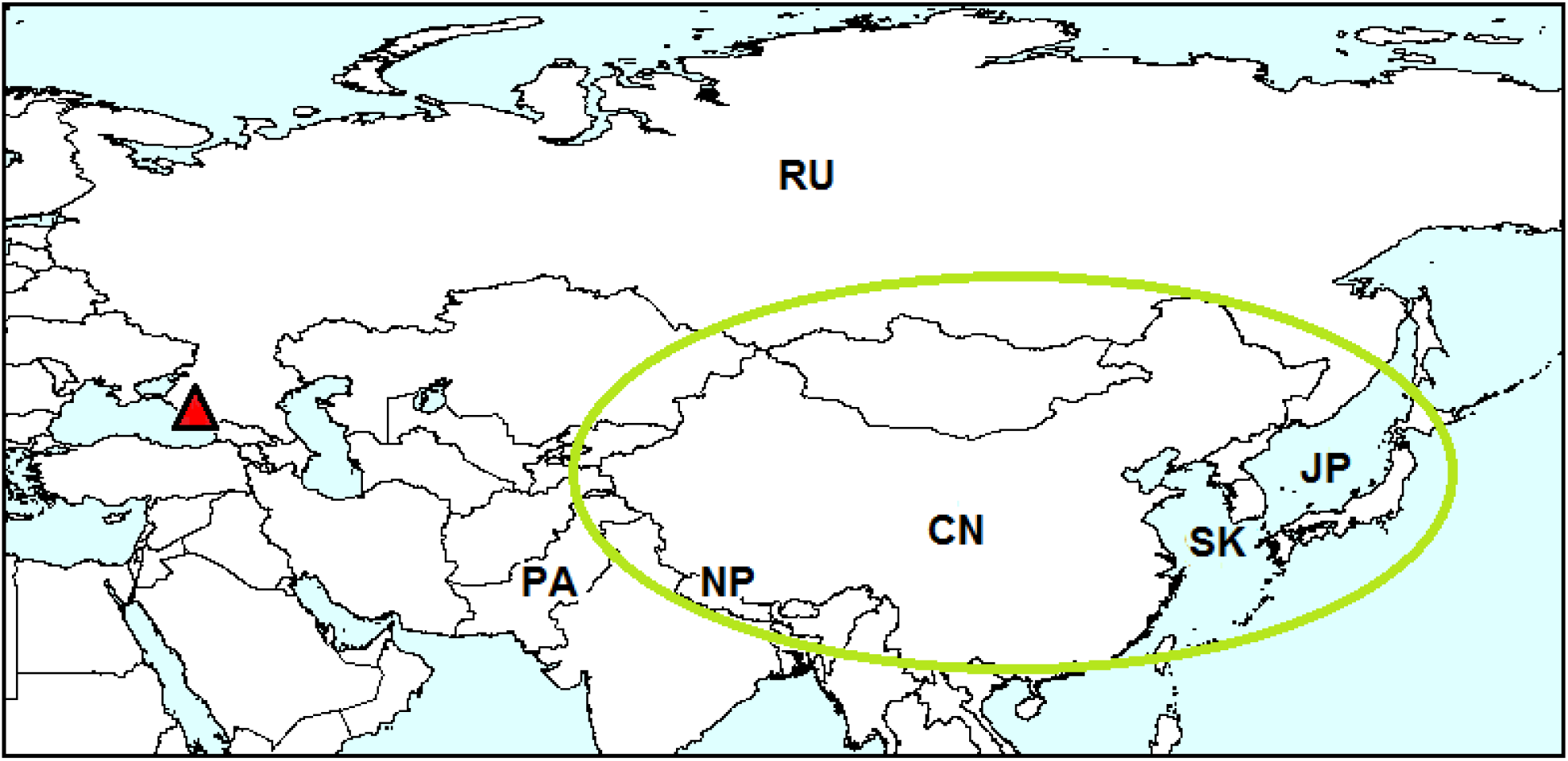
Distribution of *Medythia nigrobilineata. Red triangle* – locality of the first record outside the native range: Sochi. Native range is circled with green. CN – China, JP – Japan, NP – Nepal, PA – Pakistan, RU – Russia, SK – South Korea. Sources of information: Medvedev (1992), Beenen (2010), Warchałowski (2010), Toepfer et al. (2014), Bieńkowski and Orlova-Bienkowskaja (2018b).

#### Invasion status

Though we have found only one specimen, it is likely that it represents a population (at least temporal). First, the likelihood of collecting an individual in nature from current introduction rather than from a breeding population is vanishingly small. Second, the establishment of *M. nigrobilineata* in Krasnodar Krai is quite possible, since soybean is widely cultivated in this region (Kondratenko 2012). Special surveys of soybean plantations are necessary to reveal, if the pest is established.

#### Economic impact

Adults and larvae of *M. nigrobilineata* are known to be serious pests of soybeans in China, Japan and the Russian Far East (Ogloblin 1936; Zhanglin et al. 1997; Moseyko 2010; Takei et al. 2014). So establishment of this species in Krasnodar Region pose a serious threat to soy bean production.

### 2013 – *Chrysolina americana* (Linnaeus, 1758) (Chrysomelinae)

#### Native range

Rosemary beetle is native to Mediterranean countries: Albania, Croatia, France, Greece, Italy, Malta, Portugal, Slovenia, Spain, Serbia, Macedonia, Algeria, Morocco, Tunisia and Turkey (Fig. 5) (Beenen and Roques 2010; Beenen 2010). In particular, it occurs on islands: Mallorca, Corsica, Sardinia, Cyclades, Crete, Madeira, North Aegean Islands and Malta (Pasqual et al 2017).

**Fig. 5.**
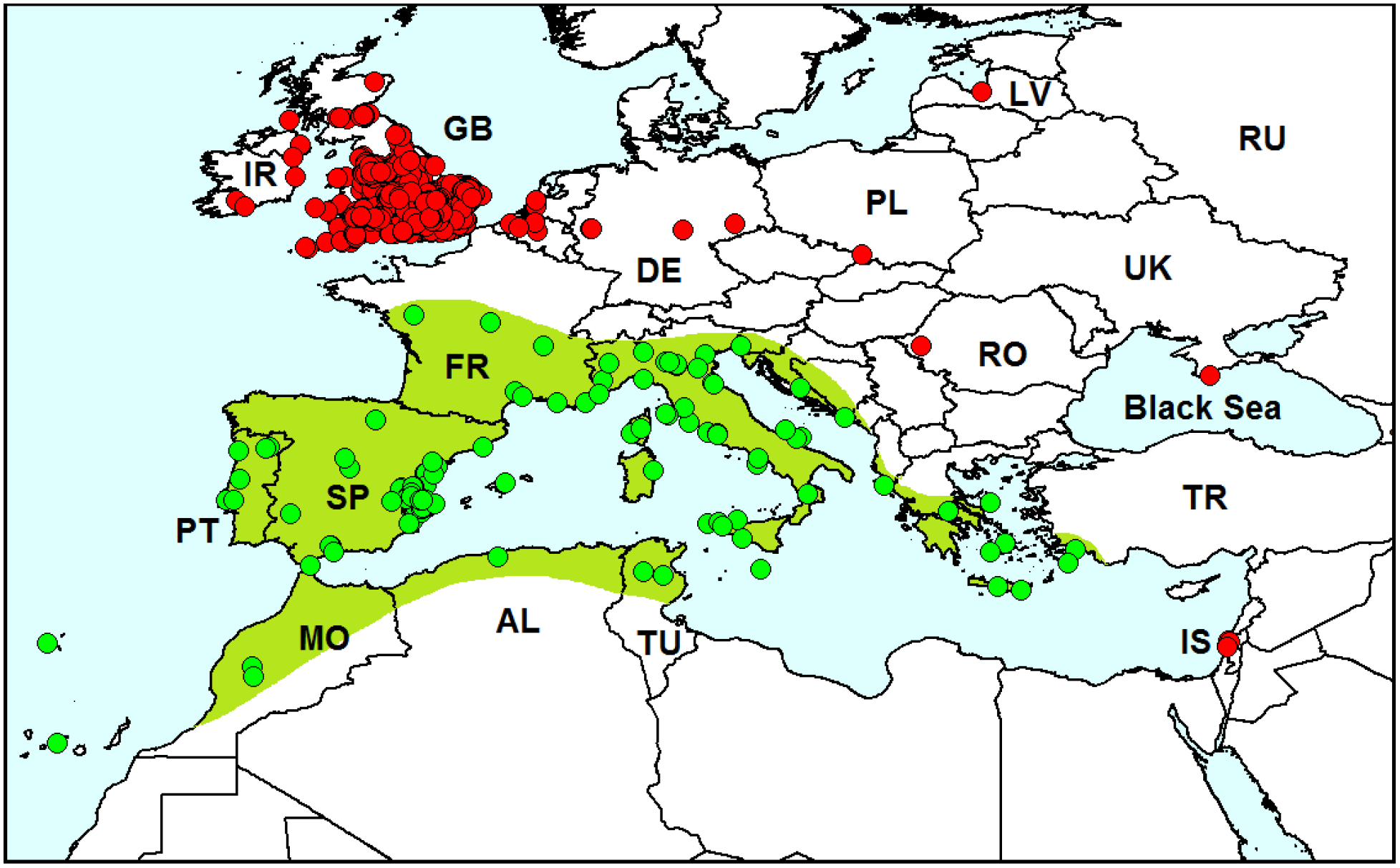
Distribution of *Chrysolina americana. Red dots* – localities, where *Ch. americana* was recorded outside its native range, *green dots* on the *green background* – localities in the native range. Sources of information: examined specimens from museum collections (AB, HNHM, MTD, NHMW, NMP, NO, RI, ZIN) and literature: Bukejs and Telnov (2010), Beenen (2010), Borowiec et al. (2011), Friedman (2016), Pasqual et al (2017), GBIF (2017b). Database on localities of *Chrysolina americana* is provided in electronic supplementary material 4.

#### Invasion history

Before the 1990s specimens of *Ch. americana* were several times collected outside the native range: in 1936 and 1938 in Belgium (GBIF 2017b), before 1950 in Romania, Austria and Germany (examined specimens from MTD) and in 1963 in the United Kingdom (Johnson 1963). But probably these findings reflect just temporary populations. When *Ch. americana* was again recorded from the United Kingdom in the 1990s, MacLeod (2002) supposed that it would not establish because of the cold climate. But surprisingly *Ch. americana* established and began to spread quickly. Now it is common all over the United Kingdom and Ireland (Royal horticultural society 2014; GBIF 2017b). It was also reported to be introduced to several other European countries and Israel (Table 1).

**Table 1.**
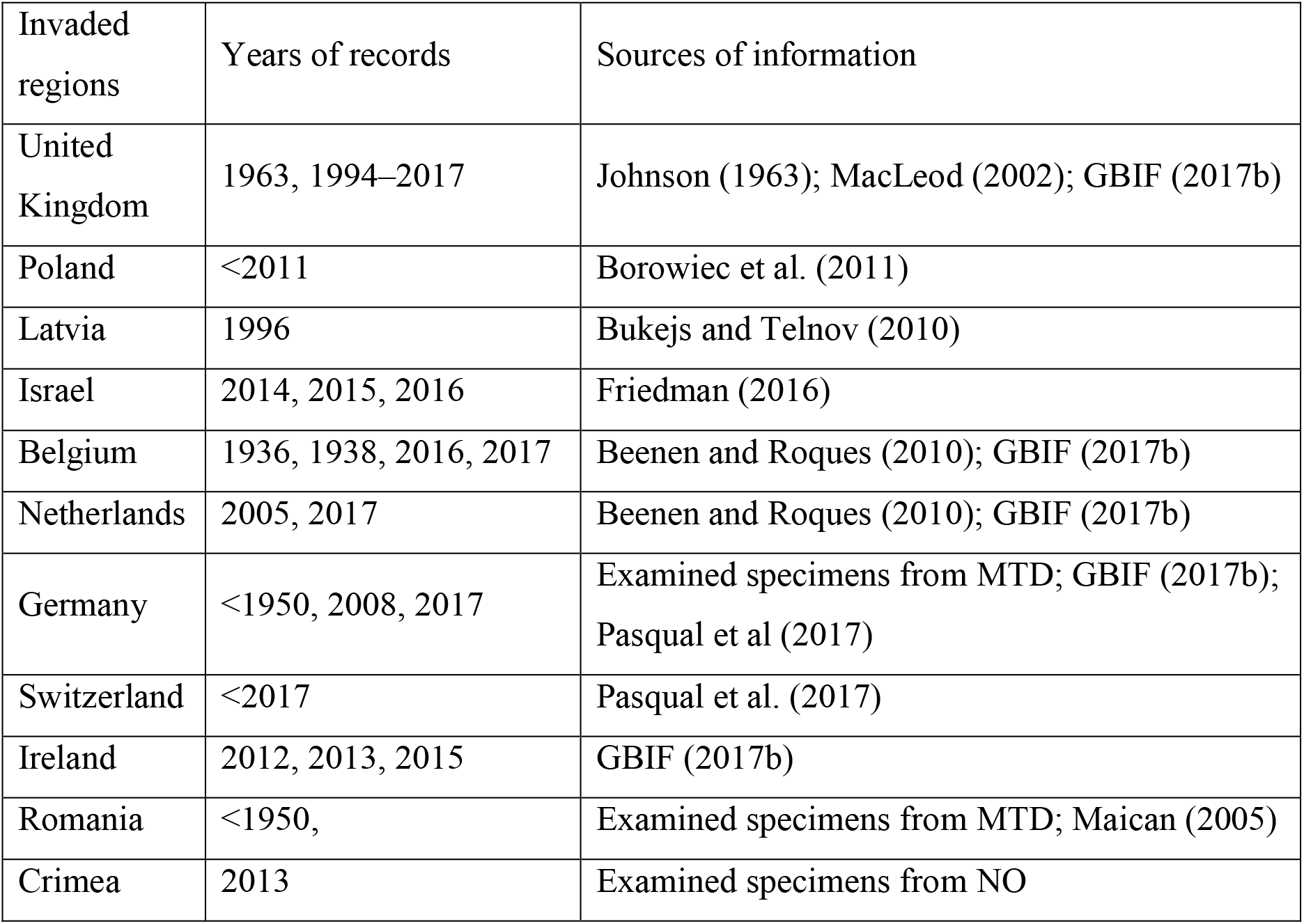
Records of *Chrysolina americana* outside its native range.

Records of *Ch. americana* from Poland, Romania, Germany and Switzerland were supposed to be questionable because of Mediterranean origin of the species (Maican 2005; Borowiec et al. 2011; Pasqual et al. 2017). We suppose that these records indicate the cases of introduction of the species outside its native range. It is difficult to determine, which records indicate establishment of species and which represent translocations or temporal populations.

#### Record in Crimea

*Chrysolina americana* has not been recorded from Russia or the Ukraine till now. Here we present the first record of *Ch. americana* on the Crimean peninsula: 13 specimens of this species were collected in Crimea in 2013 [Yalta, Bakhchisaray highway, sanatorium “Uzbekistan”, 225 m a.s.l., on *Rosmarinus officinalis*, 01.06.2013, leg. N.V. Okhrimenko].

#### Biology

Adults and larvae feed on leaves of plants of the family Lamiaceae: *Rosmarinus officinalis, Lavandula* spp., *Salvia* spp., *Thymus* spp., *Perovskia atriplicifolia* and others (Friedman 2016).

#### Vector of dispersal

Because *Rosmarinus* and *Lavandula* are popular garden plants throughout Europe, *Ch. americana* has been translocated outside its native range along with its host plants (Beenen and Roques 2010). Opinions about natural dispersal abilities of *Ch. americana* are contradicting. MacLeod (2002) states that *Ch. americana* is flightless and therefore is restricted in its dispersal abilities. But Beenen and Roques (2010) indicate that this species has good flight capacities and disperses naturally by flight. Webster et al. (2017) suppose that expansion of the rosemary beetle to the United Kingdom has been expedited by climate change.

#### Invasion status

It is not quite clear if *Ch. americana* is established in Crimea or just a temporal population was recorded. Further observations are necessary to answer this question, since example of the population in the United Kingdom shows that this species is able to be established quickly and become a serious invasive pest.

#### Economic impact

Rosemary beetle is a garden pest. It damages the foliage and flowers of various aromatic plants including lavender, rosemary and sage (Friedman 2016). These plants are widely cultivated in Crimea. So if *Ch. americana* becomes abundant, it could cause negative economic consequences in the region.

### 2013 – *Epitrix hirtipennis* (Melsheimer, 1847) (Alticinae)

#### Native range

Tobacco flea beetle is native to the south of North America, north of South America and the Caribbean Islands (Fig. 6) (Riley et al. 2003; Bieńkowski and Orlova-Bienkowskaja 2017a).

**Fig. 6.**
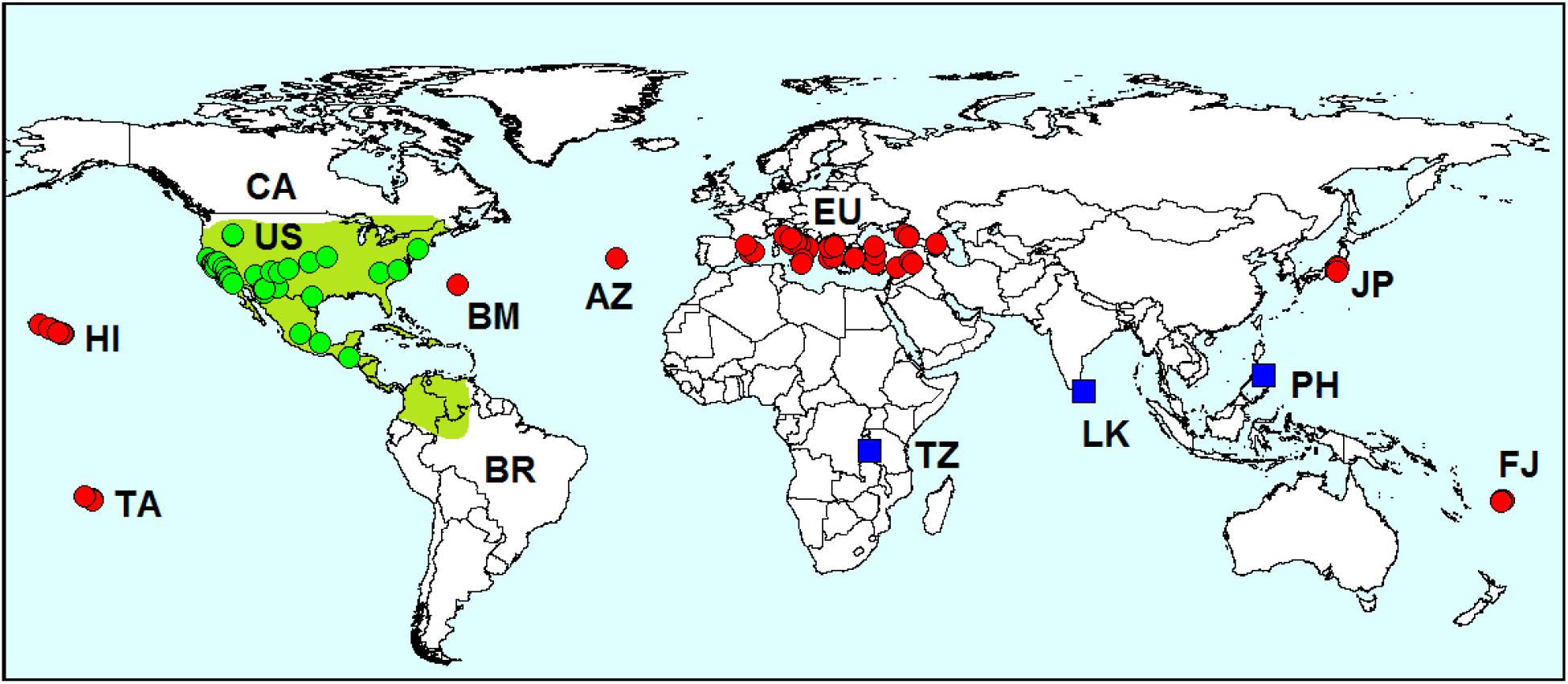
Geographical distribution of *Epitrix hirtipennis. Red dots*-localities outside the native range, *green dots* on the *green background* – localities in the native range, *blue squares* – questionable records. AZ – Azores, BM – Bermuda, BR – Brazil, CA – Canada, EU – Europe, HI – Hawaii, FJ – Fiji, JP – Japan, LK – Sri Lanka, PH – Philippines, TA – Tahiti, TZ – Tanzania, US – the United States. Sources of information about records in the native range: examined specimens from VNIIKR, ZIN and literature: Chamberlin and Tenhet (1923), Sannino et al. (1985), Furth and Savini (1996), GBIF (2017c). Sources of information about records outside the native range are indicated in Table 2. Database on localities of *Epitrix hirtipennis* is provided in electronic supplementary material 5.

**Table 2.** Records of *Epitrix hirtipennis* outside its native range

| Invaded regions | Years of records | Sources of information |
| --- | --- | --- |
| Hawaii | 1892, 1893, 1927, 1942, 1960, 1961, 1962, 1967, ~1979, ~2002 | Sharp (1900); Samuelson (1973); Gourves and Samuelson (1979); Nishida (2002), GBIF (2017c) |
| Bermuda | <1924, 1969 | Ogilvie (1924); Hilburn and Gordon (1989) |
| Tahiti | 1973, 1974, 1977, 1978 | Gourves and Samuelson (1979) |
| Fiji | 1978 | Gourves and Samuelson (1979); Waterhouse (1997); GBIF (2017c) |
| Azores | 1984 | Israelson (1985) |
| Mainland Italy | 1983, 1990 | Sannino et al. (1984); Sannino and Balbiani (1990); Deseö et al. (1993) |
| Greece | 1988, 2004 | Lykouressis (1991); Deligeorgidis et al. (2007) |
| Turkey | 1993, 2010, 2013, 2015, 2016 | Döberl (1994a); Ekiz et al. (2013); Bolu (2016); Aslan and Başar (2016); Özdikmen et al. (2017) |
| Macedonia | 1996, 2011, 2013 | Krsteska et al. (2009); Spasov et al. (2013); Krsteska and Spirkoski (2017) |
| Albania | <1997 | Krsteska and Spirkoski (2017) |
| Balearic Islands | 1998, 2016 | Petitpierre et al. (2017) |
| Bulgaria | <2000 | Trenchev and Tomov (2000); Gruev (2006) |
| Sicilia | 2000, 2010 | Baviera and Biondi (2015) |
| Syria | 2002 | Gruev and Döberl (2005) |
| Japan | 2011, 2016 | Harada and Takizawa (2012); Kim et al. (2017) |
| Russia | 2013, 2016 | Orlova-Bienkowskaja (2014) and AB |
| Georgia | 2014 | Aslan et al. (2017) |
| Croatia | <2015 | Raspudić et al. (2015) |
| Mainland Spain | 2015 | Viñolas et al. (2016) |
| Philippines (questionable record) | <1987 | Martin and Herzog (1987); Deseö et al. (1993) |
| Sri Lanka (questionable record) | <1923 | Chamberlin and Tenhet (1923); Deseö et al. (1993) |
| Tanzania (questionable record) | 2013 | Ntibiyoboka (2014) |

#### Invasion history

*Epitrix hirtipennis* began to spread outside its native range in the end of the 19th century (Figs 6, 7; Table 2). First it was introduced to islands in Atlantic and Pacific oceans: Hawaii, Bermuda, Tahiti, Fiji and Azores and has become common there.

In 1983 this species was found in Europe for the first time in Northern Italy (Sannino et al. 1985). It was the first alien flea-beetle introduced to Europe. It then spread to South and Central Italy, Greece, Turkey, Spain, Macedonia, Bulgaria, Syria, European Russia and Georgia. In 2011 it was found that *E. hirtipennis* was common on Honshu (Japan) (Harada and Takizawa 2012). In some reviews the records from Sri Lanka (Chamberlin and Tenhet 1923), Philippines (Martin and Herzog 1987; Deseö et al. 1993) and Tanzania (Ntibiyoboka 2014) are mentioned. But these records are doubtful, since no references to the source of information are given.

**Fig. 7.**
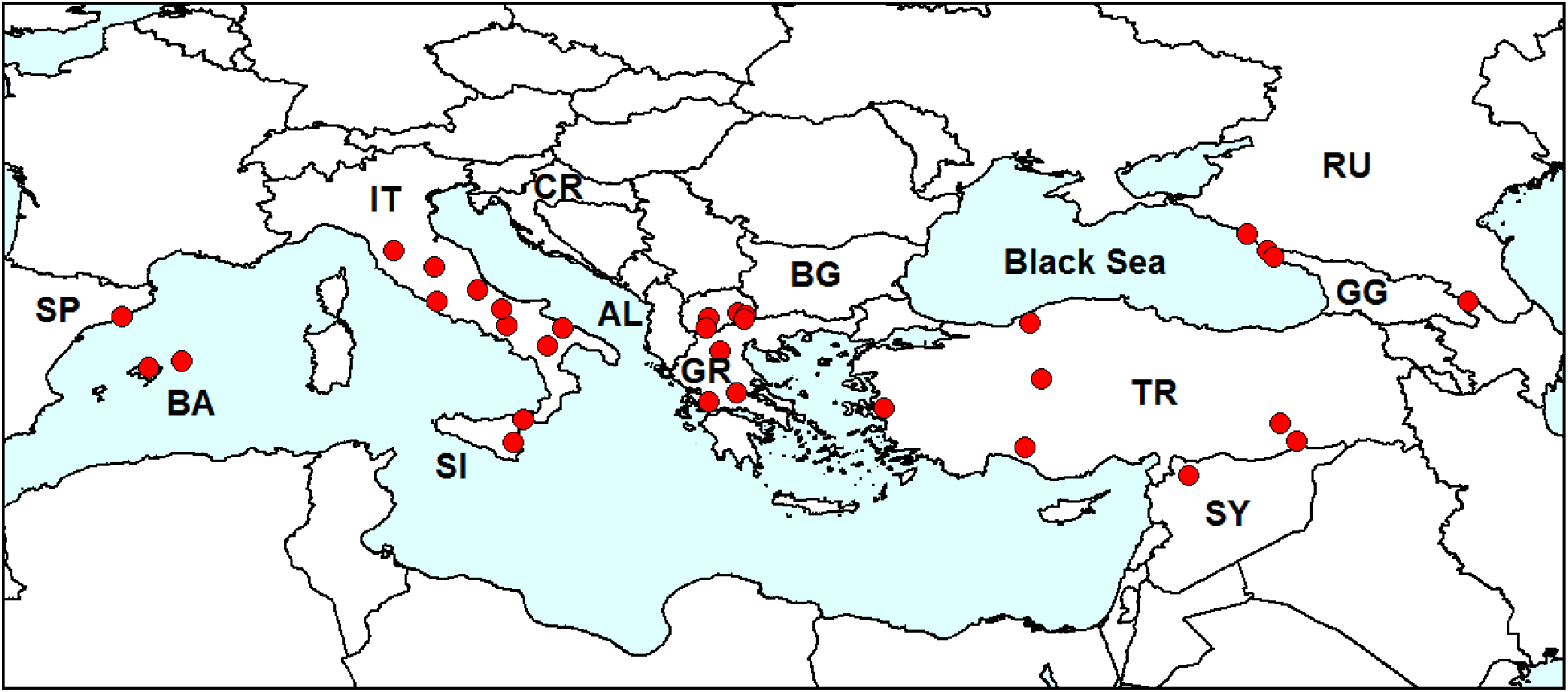
Distribution of *Epitrix hirtipennis* in Europe. *Red dots* – localities in Europe. IT – Italy (first record in 1983), GR – Greece (1988), TR – Turkey (1993), AL – Albania (<1997), BA–Balearic Islands (1998), SY – Syria (2002), RU – Russia (2013), GG – Georgia (2014), CR–Croatia (<2015), SP – mainland Spain (2015), Sources of information are indicated in Table 2. Database on localities of *Epitrix hirtipennis* is provided in electronic supplementary material 5.

#### Records in European Russia

In 2013 *E. hirtipennis* was first recorded in the Caucasus – in the cities of Sochi and Tuapse on the Black Sea coast (south of European Russia: Krasnodar Region) (Orlova-Bienkowskaja 2014). In 2016 we again found this species in the same region in the city of Adler. All specimens were sweep-netted on ruderal vegetation. It was supposed that *E. hirtipennis* was also found on tobacco field in Krasnodar (Plotnikova 2014), but the identification of species was incorrect. The expert in Chrysomelidae of the Caucasus N.V. Okhrimenko identified these specimens as *E. atropae* Foudras, 1860.

#### Biology

Adults feed on leaves of plants of the family Solanaceae. Larvae develop on roots. The review of host plants of *E. hirtipennis* in different regions was made by Bieńkowski and Orlova-Bienkowskaja (2016). In Italy *E. hirtipennis* has been observed to shift onto native Solanaceae (Beenen 2006). We collected *E. hirtipennis* in Russia on ruderal vegetation.

#### Vector of dispersal

Since *E. hirtipennis* was the first alien flea beetle introduced to Europe, its record on the continent puzzled experts in Chrysomelidae. It was assumed that *E. hirtipennis* arrived in Europe as aerial plankton with easterly trade winds blowing from the New World to Europe (Döberl 1994b). But we believe that the most likely vector of dispersal is an unintentional introduction of larvae in soil with imported planting material.

#### Invasion status

*Epitrix hirtipennis* is established in the region. Localities of its records in Europe and Asia indicate that the species is dispersing eastward.

#### Economic impact

This species is known mainly as a pest of tobacco, but can also feed on eggplant, potato, tomato and many other solanaceous plants (Capinera 2001).

#### Remark

The representatives of the genus *Epitrix* are especially prone to invasions. Five species established outside their native ranges in other continents and islands (Fig. 8). One more representative of the genus – *Epitrix setosella* (Fairmaire, 1888) was reported to be introduced outside its native range – namely from East Asia to Georgia (Aslan et al. 2017). But our examination of the specimens identified as “*E. setosella”* from collection of G.O. Japoshvili has shown that the identification is incorrect, and these specimens belong to *E. pubescens* (Koch, 1803).

**Fig. 8.**
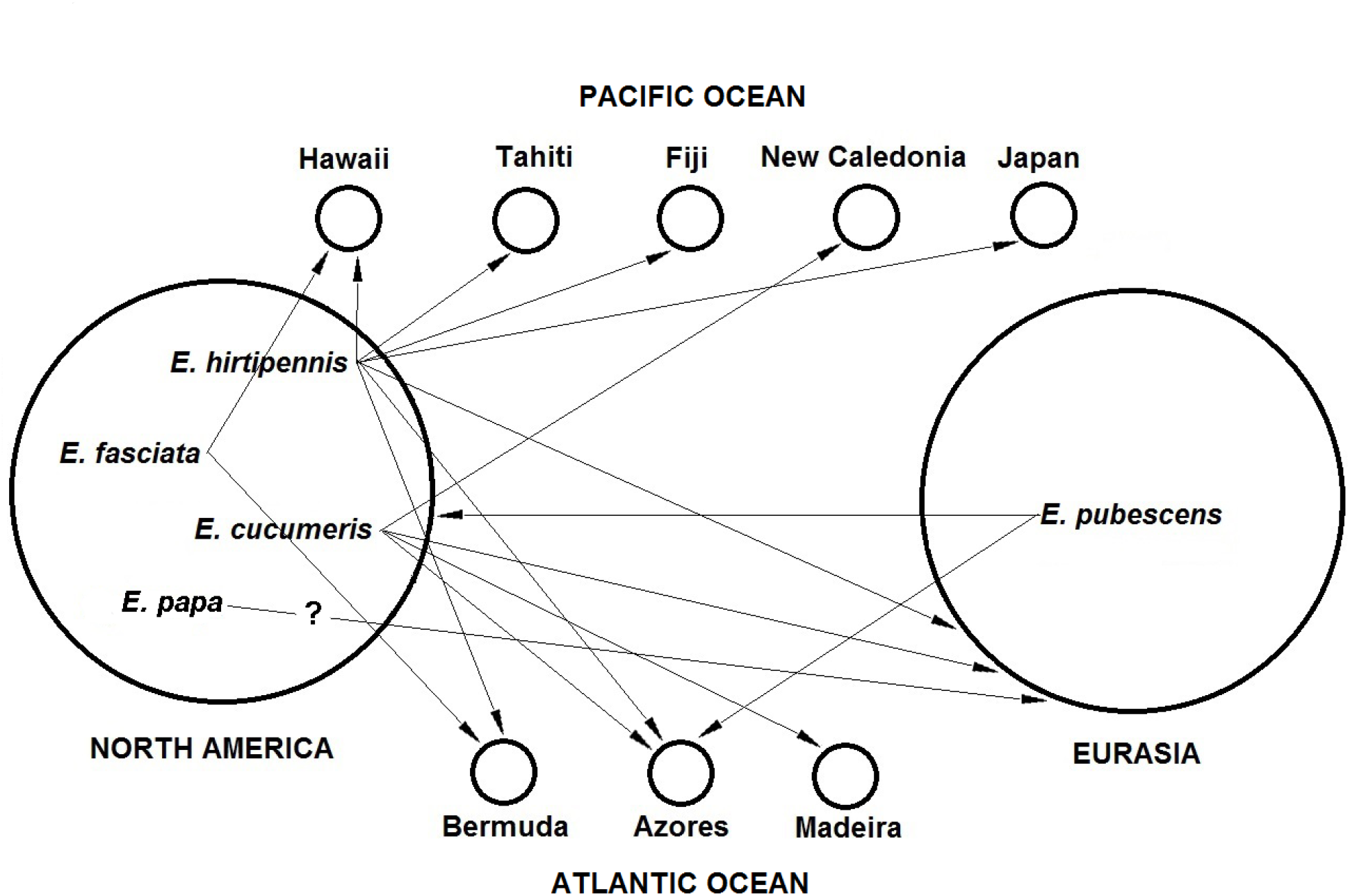
Invasions of the species of the genus *Epitrix* from their native ranges to other continents and islands in Pacific and Atlantic oceans (Bieńkowski and Orlova-Bienkowskaja 2016).

### 2011 – *Diabrotica virgifera* LeConte, 1868 (Galerucinae)

#### Native range

The western corn rootworm originates from the New World. The initial range is Mexico or Central America (Lombaert et al. 2017). Now the range in the Americas includes Canada, Costa Rica, Guatemala, Mexico, Nicaragua and the USA (EPPO 2017).

#### Invasion history

Man has greatly increased the range of this species in the Americas by the cultivation of corn (Lombaert et al. 2017). In Europe *D. virgifera* was first observed near Belgrade airport, Serbia in 1992 (Bača 1994). After several introduction events to different regions of Europe (Ciosi et al. 2008) the species has become widespread. Now it is recorded in at least 22 European countries (Fig. 8) (EPPO 2017).

**Fig. 9.**
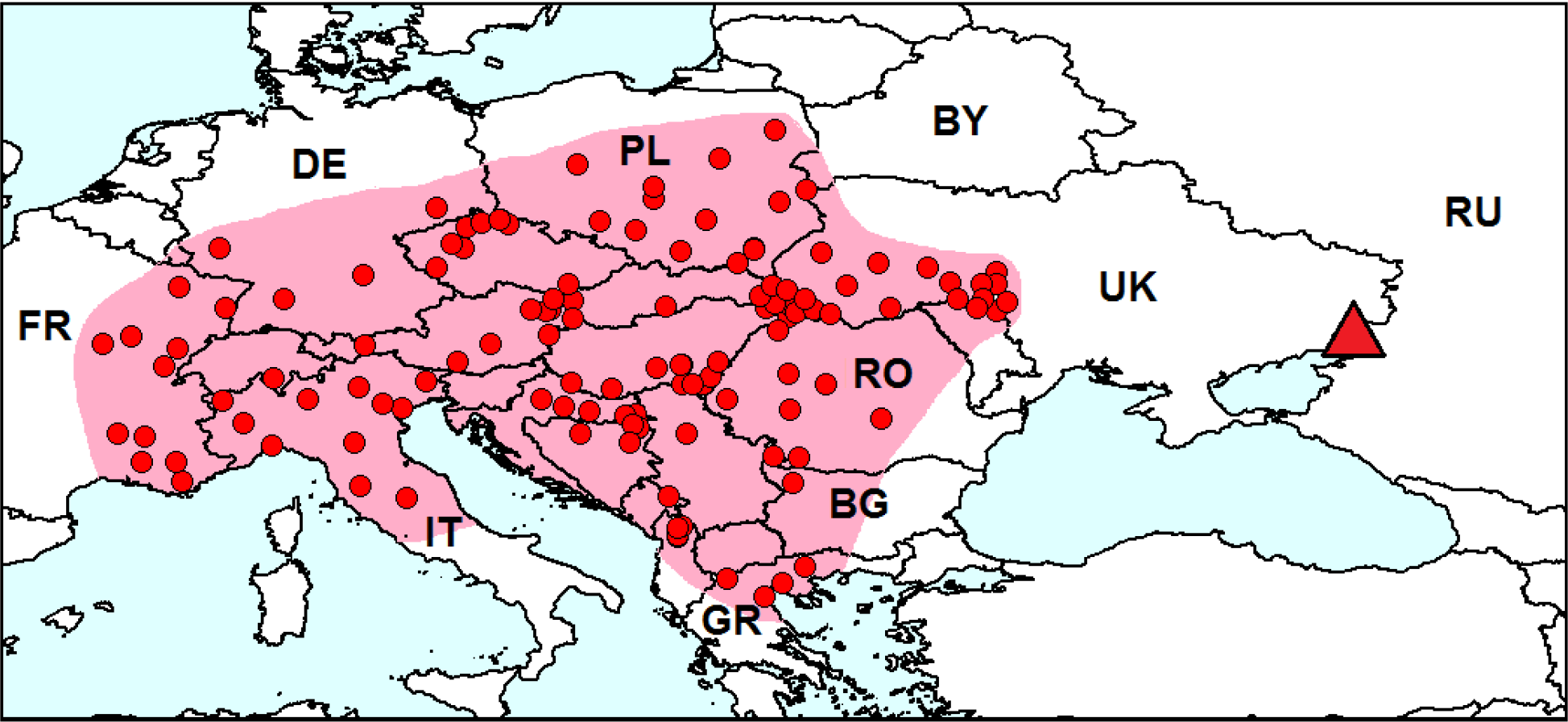
Distribution of *Diabrotica virgifera* in Europe. *Red dots* on the *pink background* – localities of species in the secondary range. *Red triangle* – locality of the interception of one specimen in Russia in 2011. BG – Bulgaria, BY – Belarus, DE – Germany, FR – France, PL-Poland, RO – Romania, RU – Russia, UK – Ukraine. Sources of information: VNIIKR (2012), Trepashko and Nadtocheva (2013), Butaliuk et al. (2016), EPPO (2017), GBIF (2017d), Manole et al. (2017). Database on localities of *Diabrotica virgifera* is provided in electronic supplementary material 6.

#### Record in European Russia

In 2011 *D. virgifera* was captured with a pheromone trap at the border of Russia in Rostov Region near the international highway (VNIIKR 2012

#### Biology

Adults of *Diabrotica virgifera* feed on leaves, silks, pollen, and young kernels of corn, larvae develop on roots (Branson and Krysan 1981).

#### Vector of dispersal

*Diabrotica virgifera* proved to have been translocated from North America to Europe several times in aircraft laden with goods and materials (Ciosi et al. 2008). The beetles fly well, so they spread in Europe both by hitchhiking and naturally (Hemerik et al. 2004).

#### Invasion status

It seems that *Diabrotica virgifera* has not been established in Russia yet, since it was intercepted only once in 2011.

#### Economic impact

*Diabrotica virgifera* is a major pest of cultivated corn. Most of the damage is caused by larvae feeding on the root system. The species is included to A2 List of quarantine pests of EPPO (2017).

### 1984 – *Phyllotreta reitteri* Heikertinger, 1911 (Alticinae)

#### Native range

We believe that the native range of *Ph. reitteri* is in Central Asia (Fig. 9), as its host plant *Lepidium latifolium* originates from this region (Hinz et al. 2008). Before the 1980s *Ph. reitteri* was recorded in Kazakhstan and Uzbekistan only (Heikertinger 1941; Lopatin 1977; Gruev and Döberl 1997). Probably the recent record from West China (Gerber et al. 2011) also belongs to the native range.

**Fig. 10.**
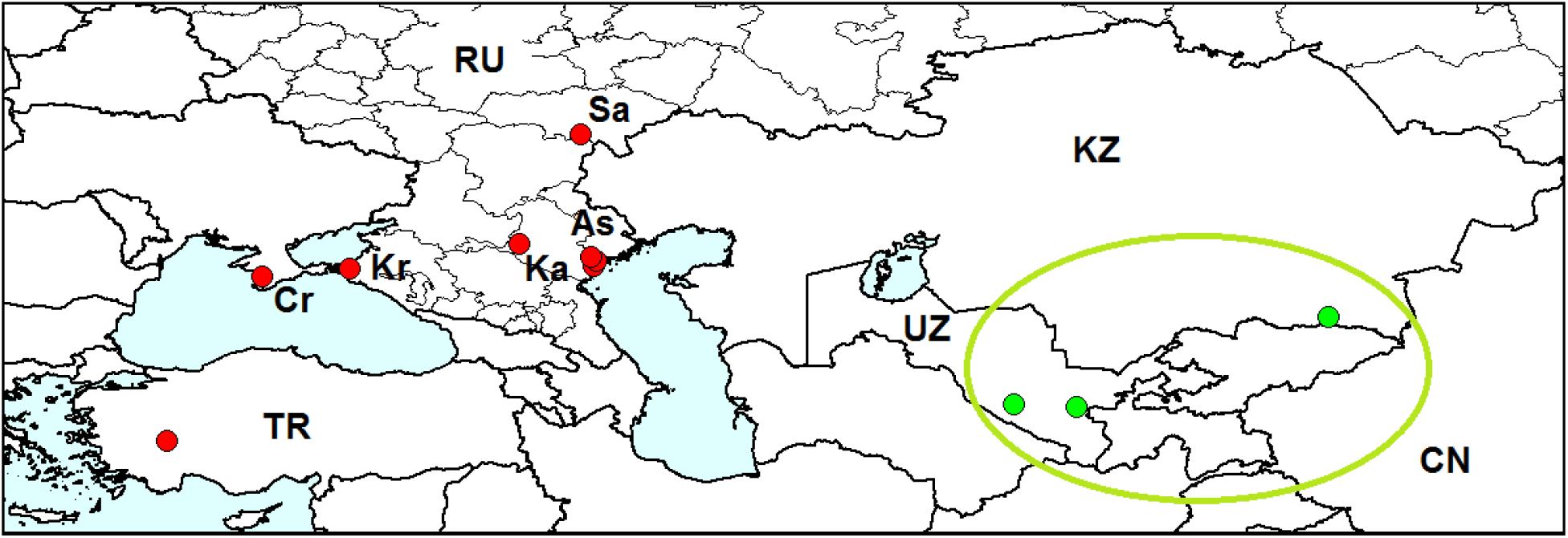
Distribution of *Phyllotreta reitteri. Green dots* localities in the native range, *red dots* – localities in the native range, red dots – localities outside the native range (records after 1984), CN – China, KZ – Kazakhstan, RU – Russia, TR – Turkey, UZ – Uzbekistan, As – Astrakhan Region, Cr – Crimea, Kr – Krasnodar Territory, Ka – Kalmykia, Sa – Saratov Region. Sources of information are indicated in Table 3.

#### Invasion history

In 1984 *Ph. reitteri* was first found outside its native range, namely in Crimea (Mosyakin 1987). Then in 1986.2012 it was found in four regions in the South of European Russia (Table 3) and in 2002 in Turkey (Gök et al. 2002).

**Table 3.** Records of *Phyllotreta reitteri* outside its native range.

| Invaded regions | Years of | Sources of information |
| --- | --- | --- |

|  | records |  |
| --- | --- | --- |
| Crimea: Simferopol | 1984 | Mosyakin (1987) and examined specimens from SM |
| Kalmykia: Elista, Lagan and Dzhalykovo | 1986, 2011, 2012 | Our collection and examined specimens from GK |
| Astrakhan Region: Liman | 2010 | Our collection |
| Saratov Region: D'yakovka | 2004 | Examined specimens from AU |
| Krasnodar Territory: Temryuk and Golubitskaya | 2007, 2008, 2010 | Examined specimens from ZIN (collected by B.A. Korotyaev and A.G. Moseyko) |
| Turkey: Denizli | 2002 | Gök et al. (2002) |

#### Distribution in European Russia

*Phyllotreta reitteri* occurs in Kalmykia, Saratov Region, Astrakhan Region and Krasnodar Territory (see Table 3).

#### Biology

Experiments carried out in cage and in field have shown that the main host plant of *Phyllotreta reitteri* is perennial pepperweed *Lepidium latifolium* (Gerber et al. 2012). Adults feed on leaves, larvae mine in petioles and shoots. This plant of Asian origin has been cultivated as a spice and vegetable since the twelfth century (Hegi 1986). Now it is widespread in Asia and Europe and has been recorded in all continents and has become an invasive weed in North America (Gerber et al. 2012). In European Russia *Ph. reitteri* occurs in moist habitats: on the banks of rivers and ponds and in irrigated parks.

#### Vector of dispersal

Unknown.

#### Invasion status

We believe that *Ph. reitteri* is alien in European Russia. It corresponds to at least four criteria of alien beetle species (Orlova-Bienkowskaja 2016) as follows:

(1) Detection of an established population of the species, which has not been recorded earlier. – *Phyllotreta reitteri* appeared outside its historically known range (Central Asia) in the 1980s.
(2) Disjunction of the range, which cannot be explained by disjunction of suitable landscapes or host plant ranges. – The range of *Ph. reitteri* consists of two parts. The first one is in Central Asia, and the second one is in the south of European Russia, Crimean peninsula and Turkey. The distance between these two parts is more than 1500 km.
(3) Expansion of a part of the range isolated from its main part. – Records in Turkey, Astrakhan and Saratov Regions in the 2000s could probably indicate the expansion of the range.
(4) Feeding on an alien host plant. – *Lepidium latifolium* originates from Central Asia (Hinz et al. 2008).

It is unlikely, that the species occurred in the region, but stayed unnoticed for a long time, because this beetle is large for the genus *Phyllotreta* and has characteristic bright coloration. There are no specimens of *Phyllotreta reitteri* collected in the South of European Russia in the rich collection of Zoological Institute of Russian Academy of Sciences, though there are thousands specimens of other *Phyllotreta* species collected in this region in the end of the 19th and the first half of the 20th century.

#### Economic impact

*Phyllotreta reitteri* is regarded as a potential biological control agent of perennial pepperweed (Hinz et al. 2008).

### 1982 – *Zygogramma suturalis* (Fabricius, 1785) (Chrysomelinae)

#### Native range

Ragweed leaf beetle is native to the USA and the south of Canada (Riley et al. 2003).

#### Invasion history

*Zygogramma suturalis* (Fig. 10) was introduced to the USSR from Canada and the USA for control of one of the most noxious invasive weeds *Ambrosia artemisiifolia* (Kovalev et al. 2015). The beetles were released in 16 provinces of the USSR: in European Russia, Ukraine, Georgia, Kazakhstan and the Far East. The most intensive work was performed in the south of European Russia, mainly in Rostov Region and in Stavropol and Krasnodar Territories (Kovalev et al. 2015). First release (1500 specimens) was made in the vicinity of Stavropol in 1978. In 1981–1983 *Z. suturalis* became abundant and began to spread quickly. Now it is rather widespread in the south of European Russia and occurs also in southeast of the Ukraine and in Georgia (Sergeev 2012 and personal communication by M.E. Sergeev). In 1985 one specimen was found in Turkey, but the species did not establish, since there were no other records (Aslan et al. 2003; Özdikmen et al. 2014 and personal communication by H. Özdikmen).

**Fig. 11.**
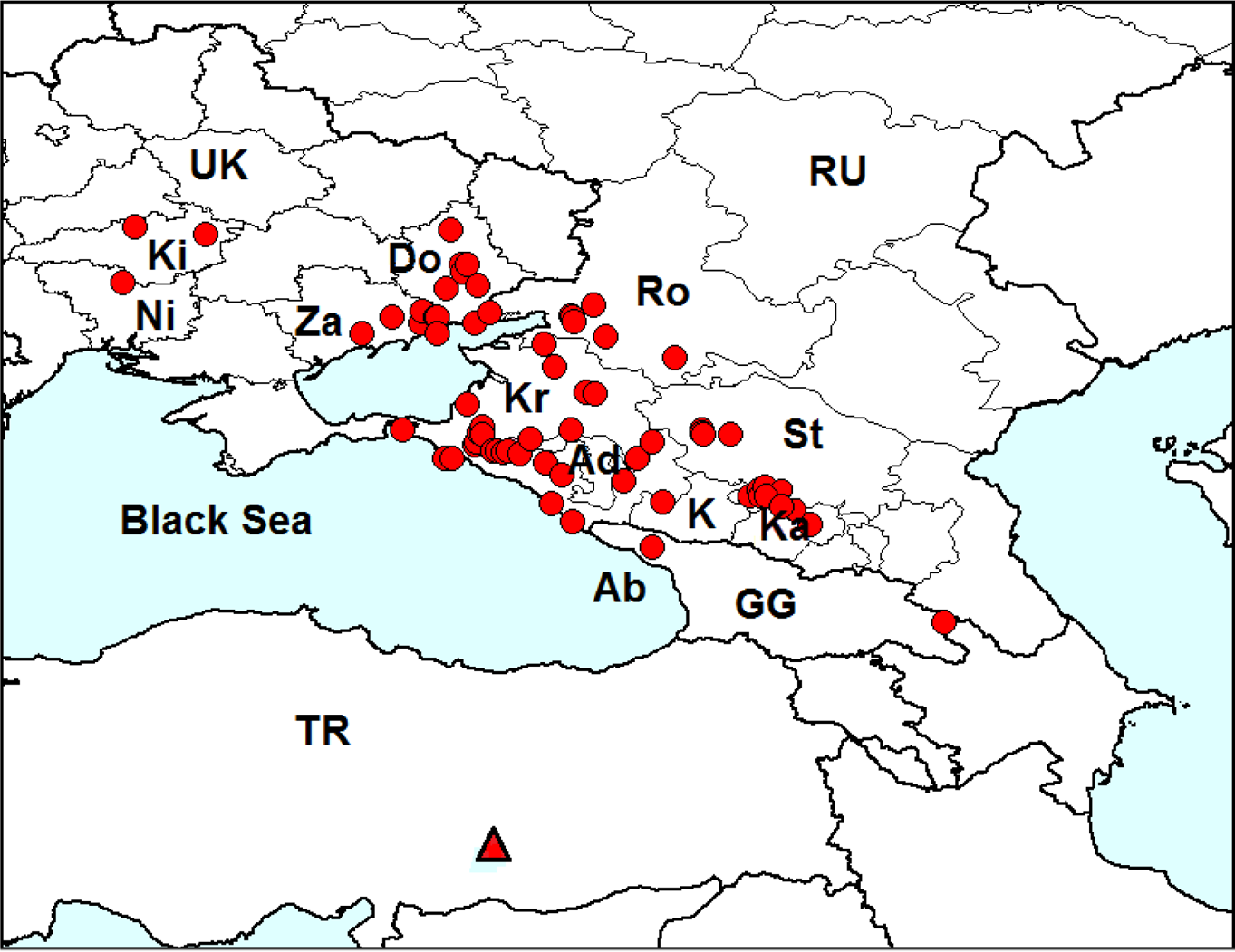
*Zygogramma suturalis*. Distribution of *Zygogramma suturalis* in Europe. *Red dots*-localities of species. GG – Georgia, RU – Russia, TR – Turkey, UK – Ukraine, Ab – Abkhazia, Ad – Adygea, Do – Donetsk Region, K – Karachay-Cherkessia, Ka – Kabardino-Balkaria, Ki-Kirovograd Region, Kr – Krasnodar Territory, Ni – Nikolaev Region, Ro – Rostov Region, St-Stavropol Territory, Za – Zaporizhje Region. Sources of information: examined specimens from the following collections: AB, AK, BSU, PR, TM, VNIIKR, YK and literature: Yaroshenko (1994), Aslan et al. (2003), Reznik and Spasskaya (2005), Sergeev (2008, 2010, 2011, 2013). Database on localities of *Zygogramma suturalis* is provided in electronic supplementary material 7.

*Zygogramma suturalis* when introduced to the south of European Russia, showed rapid evolutionary changes in flight capacity (development of flight ability and morphological changes) within only five generations (Kovalev 2002). The morphological changes were so significant, that the new subspecies *Zygogramma suturalis volatus* Kovalev, 2002 was described (Kovalev 2002).

In 1982–1985 beetles from Stavropol Territory were released in Primorje (Far East). In the 1990s it was supposed that the population of *Z. suturalis* in the Far East disappeared. But in 2010 it was discovered that this species still occurred there, though it was not abundant (Aistova and Bezborodov 2015). *Zygogramma suturalis* was also released in Croatia and Australia, but failed to establish, releases of *Z. suturalis* in China in 1985 resulted in establishment in some locations (Gerber et al. 2011). Record of *Z. suturalis* from Kazakhstan (Gerber et al. 2011) probably refers to the releases rather than to established populations. There are no current records of *Z. suturalis* from this country (Lopatin 2010).

#### Distribution in European Russia

Now *Z. suturalis* occurs in five provinces in the south of European Russia: Stavropol Territory, Krasnodar Territory, Adygea, Rostov Region, Karachay-Cherkessia (Kovalev et al. 2013; Reznik and Spasskaya 2005) and Kabardino-Balkaria (unpublished data by T.A. Mogilevich).

#### Biology

Larvae and adults feed on leaves, shoots and inflorescences of *Ambrosia artemisiifolia* (Kovalev and Medvedev 1983). The species is very abundant in some localities, occurs in river valleys, on the fringes of forests, and in saline areas (Sergeev 2013).

#### Vector of dispersal

Intentional introduction for biological control of *Ambrosia artemisiifolia*.

#### Invasion status

*Zygogramma suturalis* is established in European Russia.

#### Economic impact

Soon after the introduction of *Z. suturalis* to the Caucasus the density of some populations was as high as 100 million specimens per km^2^. *Zygogramma suturalis* completely destroyed *Ambrosia* in some locations. But 10 years after the introduction surveys revealed that in spite *Z. suturalis* was still abundant in some places, however it does not significantly affect the general density of the host plant. The system “plant-phytophagous” reached the equilibrium (Kovalev et al. 2013; Reznik et al. 2008). The same situation is observed in the Far East (Aistova and Bezborodov 2015).

### 1958 – *Leptinotarsa decemlineata* (Say, 1824) (Chrysomelinae)

#### Native range

The Colorado potato beetle originates from central highlands of Mexico (Alyokhin et al. 2013).

#### Invasion history

From the beginning of the 19th to the beginning of the 20th century *L. decemlineata* colonized whole North America (Ivanchik and Izhevsky 1981; Alyokhin et al. 2013). In 1922 *L. decemlineata* was found to be established in Bordeaux (France) and began to spread in Europe. Now it is distributed almost throughout Europe and northern Asia (Kippenberg 2010; Maslyakov and Izhevsky 2011; Alyokhin et al. 2013; EPPO 2017). Presently, the Colorado potato beetle damages potato crops all over Europe, Asia Minor, Iran, Central Asia, and western China (Alyokhin et al. 2013). General distribution of *Leptinotarsa decemlineata* is well known (EPPO 2017), so we present the map of distribution of this species in Russia only (Fig. 11).

**Fig. 12.**
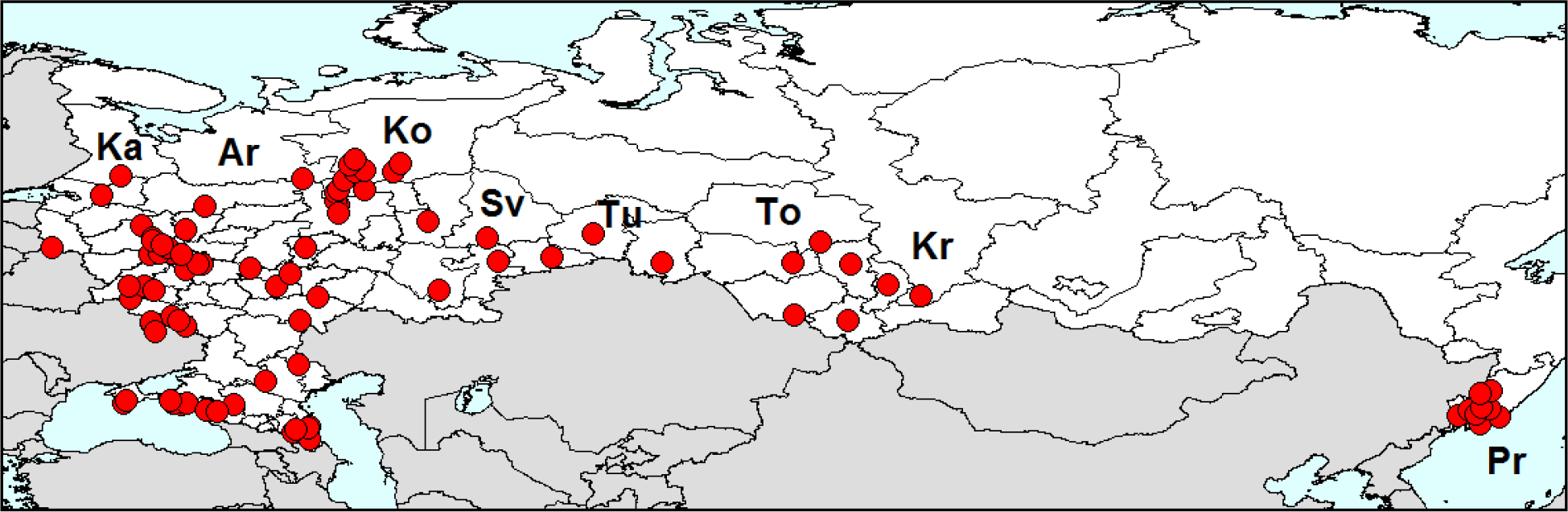
Distribution of *Leptinotarsa decemlineata* in Russia. Ar – Arkhangelsk region, Ka-Karelia, Ko – Republic of Komi, Kr – Krasnoyarsk Territory, Pr – Primorsky Territory. To-Tomsk Region, Tu – Tumen Region, Sources of information: examined specimens from the following collections: AB, AP, AR, BNR, BSU, EI, GK, MD, NN, OSU, PP, VF, VNIIKR, ZMMU, and literature: Matsishina (2011), Guskova (2013), Popova (2014). Database on localities of *Leptinotarsa decemlineata* is provided in electronic supplementary material 8.

#### Expansion to Russia

In 1958 expansion of the pest range reached the western border of the USSR (Maslyakov and Izhevsky 2011). Now the Colorado potato beetle is common all over European Russia, even in the north: in Leningrad Region (Maslyakov and Izhevsky 2011) and the Republic of Komi (Dolgin and Bieńkowski 2011). Its range has expanded to the most part of Siberia. The northern boundary of the area passes through Karelia, Arkhangelsk Region, Republic of Komi, Tumen Region, Tomsk Region and Krasnoyarsk Territory. Since 2000 the isolated part of the range exists also in the Far East – in Primorsky Territory (Maslyakov and Izhevsky 2011).

#### Biology

In European Russia *L. decemlineata* feeds on *Solanum tuberosum, S. lycopersicum, S. melongena, S. laciniatum, S. dulcamara, Hyoscyamus niger* and *Atropa belladonna* (Medvedev and Roginskaya 1988; Maslyakov and Izhevsky 2011; Bienkowski and Orlova-Bienkowskaja 2015). It occurs not only in agricultural landscapes, but also in undisturbed natural communities, in particular, on river banks (Bieńkowski 2011).

#### Vector of dispersal

This pest enters new territories mainly because of unintentional transportation with potato and self-dispersal of beetles sometimes aided with the winds (Ivanchik and Izhevsky 1981).

#### Economic impact

*Leptinotarsa decemlineata* is the most devastating pest of potato and other cultivated plants of the family Solanaceae (Alyokhin et al. 2013).

### Species established before the 20th century – archeoinvaders

It can be assumed that some leaf beetles species spread from their native ranges with the spread of agricultural plants and weeds centuries ago. In particular, Beenen (2006) argued that the combination of *Buglossoides arvensis* and *Longitarsus fuscoaeneus* Redtenbacher 1849 might have taken the route from southwest Asia where they spread with agriculture to large parts of the temperate parts of the Northern hemisphere. By analogy with the term archeophytes, which is applied to alien plants, the term “archeoinvaders” was introduced for animals (Khlyap et al. 2010). It is very difficult to distinguish such established alien species from native ones. Thus, a number of species which are at present considered as native may indeed be originally alien (Beenen and Roques 2010). In some cases detail analysis of ecology and long-term dynamics of ranges could help to reveal archeoinvaders by the criteria similar to those used by botanists for revealing of archaeophytes (Orlova-Bienkowskaja 2016).

In particular, *Chrysolina eurina* (Frivaldszky, 1883) (Chrysomelinae) was argued to be probably an archeoinvader in Europe (Orlova-Bienkowskaja 2013a). The detailed analysis of all localities of this species available from literature and museum collections has shown that its range is disjunctive: consists of three isolated parts, which are at a great distance from each other. The western part is situated in Central Europe (Austria, Hungary, Slovakia, Romania, Poland, Czech Republic), the middle one – in European Russia (Moscow Region, Tambov Region, Nizhny Novgorod Region, Penza Region, Vladimir Region, Samara Region and Ryazan Region), and the eastern part – in the south-east of West Siberia (Kemerovo Region, Republic of Khakassiya, Krasnoyarsk Territory, Altai Republic). We believe that western and the middle parts probably appeared as a result of invasions more than 100 years ago. Firstly, the only host plant of this leaf-beetle is *Tanacetum vulgare*, a weed, which was introduced to Europe in antiquity (Pyšek et al. 2002). Secondly, *Ch. eurina* in Europe occurs mainly in the ruderal habitats, which is typical for nonnative species. Thirdly, the unintentional introduction of beetles by transport is quite possible, since the local populations often occur at the roadsides of highways. Disjunction of the range cannot be explained by disjunction of host plant range or suitable landscapes, since *T. vulgare* has a continuous range in Europe, and *Ch. eurina* occurs not in mountains or some kind of other rare or relict landscape, but on the contrary – in anthropogenic landscapes in highly populated areas.

Another example is *Lilioceris lilii* (Scopoli, 1763) (Criocerinae). We studied dynamics of its range and came to conclusion that the species could be introduced from Asia to Europe in the 16th–17th century with bulbs of lilies (Orlova-Bienkowskaja 2013b, 2016). *Lilioceris lilii* meets several criteria of alien species: (1) Until the beginning of the 20th century *L. lilii* had a disjunctive range consisting of the Asian and the West European subranges separated by a vast territory where the species was not found. The absence of *L. lilii* in collections from the greatest part of European Russia and Western Siberia cannot be attributed to insufficient material, since the distribution map of the closely related species *Lilioceris merdigera* (Linnaeus, 1758) compiled from the same sources reveals a continuous range of the latter species in the 19th century (the method of comparison of ranges of related species; see Gorodkov, 1981). Since the beginning of the 20th century, the European range of the species has been expanding in the northern and eastern directions. (2) *Lilioceris lilii* is prone to invasions, in particular, it became a common and widespread species on the British Isles and in North America where it appeared in the 1940s (Majka and LeSage 2008). (3) The main host plants of the species are cultivated *Fritillaria* and *Lilium* (Medvedev and Roginskaya 1988). (4) In Europe, *L. lilii* can be found almost exclusively in anthropogenic biotopes: gardens and flowerbeds. Despite being common, the species was only sporadically recorded outside the artificially planted areas. (5) The genus *Lilioceris* as a whole is of Asian origin, and the great majority of its species occur in Asia (Berti and Rapilly 1976). (6) The known vector of invasion of *L. lilii* to new territories is shipment of lily bulbs with soil.

Of course, it is not easy to distinguish alien beetle species established before the 20th century from native ones. All criteria listed above are mere indirect evidences of the alien status of a species in the given territory, because numerous exceptions exist. Usually it is impossible to recognize an alien species by a single criterion, but matching several criteria characterizes the species as an alien one with high probability.

## Discussion

Analysis of invasions of leaf beetles to European Russia has revealed some general tendencies. First of all, it is obvious that recent translocations and establishment of Chrysomelidae species outside their native ranges have become much more common than previously. Only three alien leaf beetle species were recorded in European Russia in the 20th century: *Leptinotarsa decemlineata, Phyllotreta reitteri* and intentionally introduced *Zygogramma suturalis*. But in 2000–2017 as much as six alien species were found in the region. It corresponds to the general tendency exponential increase in the rate of invasions of leaf beetles to Europe (Beenen and Roques 2010) and increase of invasions of beetles feeding on living plants (Orlova-Bienkowskaja 2017). It reflects the increase of international trade, especially trade of living plants and the increase of air transport. In most cases leaf beetles are translocated to the remote regions with imported plants or (as in the case of *Diabrotica virgifera)* by hitchhiking in airplanes (Hemerik et al. 2004; Ciosi et al. 2008). Most of alien leaf beetles are associated with agricultural plants. But some species (for example, *Phyllotreta reitteri)* develop on weeds.

After the establishment outside their native ranges leaf beetles can quickly spread by unintentional introduction by man and natural dispersal. For example, *Diabrotica virgifera* was first recorded in Europe in 1992, but now it has spread to at least 21 countries and become major pest of corn in some regions (EPPO 2017). *Luperomprpha xanthodera* has occupied the whole Europe from Spain to Russia in just 15 years. Therefore, the records of new alien pest leaf beetles are very important and should be published quickly.

In some cases invasions of leaf-beetles are unpredictable. For example, establishment of *Leptomona erythrocephala* native to Spain in the Caucasus is rather surprising. But in other cases emergence of alien species is easy to predict. For example, it was obvious that *Epitrix hirtipennis* could appear in the south of European Russia, since its range in Europe was expanding to the east. Similarly in the early 20th century it was obvious that *Leptinotarsa decemlineata* would appear in Europe, because of its outbreak and quick range expansion in North America. In general, if the range of some leaf beetle species is quickly expanding, or if the species has been recorded established somewhere outside its native range, this species should be regarded as a potential invader all over the world. For example, global invasion of *Epitrix hirtipennis* began from its establishment in Hawaii in the end of the 19th century (Sharp 1900).

Leaf beetles could be potentially translocated from any part of the world. Four species alien to European Russia originate from North America, two from Mediterranean region and three from Asia. Recently the invasions of beetles East Asia are becoming more important (Orlova-Bienkowskaja 2017). Establishment and quick spread of *Luperomprpha xanthodera* in Europe reflects this current tendency.

The Black Sea region is more prone to invasions of leaf beetles than other regions of European Russia. The large number of records of alien species in these regions could not be explained by more intensive survey, since we have the database of localities of leaf-beetles collected in all regions of European Russia (about 30000 localities) (Bienkowski and Orlova-Bienkowskaja 2017b). The large number of invasions in the Black Sea region corresponds to the general tendency of large number of invasions of leaf beetles in territories with warm and wet climate. For example, Italy occupies the first place in Europe on number of alien leaf beetles (Beenen and Roques 2010).

But it is difficult (or even impossible) to predict, if the introduced species could become established on the base of simple comparison of climate in its native range and invaded range, because establishment of species depends on a complex of interacting abiotic, biotic and anthropogenic factors. For example, when *Ch. americana* was for the second time recorded from the United Kingdom in the 1990s, MacLeod (2002) supposed that it would not establish because of the cold climate. But it has established and become a common pest.

Alien leaf beetles can spread to native communities and become naturalized. For example, *Zygogramma suturalis* occurs even in native undisturbed communities of nature reserves (Sergeev 2008, 2011). Usually alien leaf beetles remain strictly related to their original, alien plants (Beenen 2006; Beenen and Roques 2010). This is also true for Chrysomelidae of European Russia. But some species can feed also on native plants or cultivated plants from other regions. In particular, *Leptinotarsa decemlineata* feeds not only on cultivated plants, but also on *S. dulcamara* and *Hyoscyamus niger*. If the species is established in native communities and feed on native plants it is fully naturalized, i.e. ecologically undistinguishable from native species. Just because of the full naturalization it is difficult to distinguish between native species and species established before the 20th century.

Since the invasions of leaf beetles can cause tremendous economic consequences and since the exponential increase in the rate of invasions is recently observed, special attention should be paid to the study of these invasions. The monitoring is necessary to reveal the cases of emergence of species outside their native ranges. General trends of invasions of leaf-beetles should be examined and analyzed carefully.

## Acknowledgements

We would like to thank N.V. Okhrimenko, R.N. Ishin, G.O. Japoshvili, A.G. Koval, T.A. Mogilevich, P.V. Romantsov, Y.N. Kovalenko, P.N. Petrov, A.I. Prikhodko, A.B. Ruchin, G.A. Korostov, V.I. Filippov, E.V. Iljina, N.E. Nikolaeva and M.M. Dolgin for the possibility to study specimens from their collections, to S.Ya. Reznik, V.V. Martynov, M.E. Sergeev, S.A. Mosyakin, A.G. Moseyko, H. Özdikmen for valuable information on leaf beetle distribution and to A.S. Konstantinov for valuable remarks on the manuscript. The study was supported by Russian Science Foundation, Project No 16-14-10031.

## Notes

**Disclosure statement:** No potential conflict of interest was reported by the authors.

**Funding:** This work was supported by Russian Science Foundation under Grant 16-14-10031.

## References

Aberlenc HP (1987) Coléoptères de l’Ardèche. Premier supplément à l’inventaire de J. Balazuc (1984). Bull Mens Soc Linnéenne Lyon 56(10):320–349

Aberlenc HP (2010) Liste commentée des insectes du Bois de Païolive (Gard & Ardèche). Version du1er octobre 2010. 54 pp http://www.aberlentomo.fr/ Accessed 13 January 2018

Aistova EV, Bezborodov VG (2015) Results of the introduction of the ragweed leaf beetle Zygogramma suturalis (Coleoptera: Chrysomelidae) in Primorskii Krai. A.I. Kurentsov’s Annual Memorial Meetings 26:144–149 (in Russian)

Alyokhin A, Udalov M, Benkovskaya G (2013) The Colorado potato beetle. In: Giordanengo P, Vincent Ch, Alyokhin A (ed) Insect Pests of Potato. Global Perspectives on Biology and Management. Academic Press is an imprint of Elsevier, pp 11–29 http://dx.doi.org/10.1016/B978-0-12-386895-4.00001-6

Aslan I, Gruev BA, Özbek H (2003) A preliminary review of the Subfamily Chrysomelinae (Coleoptera, Chrysomelidae) of Turkey. Linzer Biol Beitr 35(1):581–605

Aslan EG, Başar M (2016) Flea beetles collected from olive trees of Antalya Province, including the first record of the monotypic genus *Lythraria* Bedel, 1897 (Coleoptera: Chrysomelidae) for Turkey. Turkish J Entomol 40(3):243–248. http://dx.doi.org/10.16970/ted.00746

Aslan EG, Mumladze L, Japoshvili G (2017) List of leaf beetles (Coleoptera: Chrysomelidae) from Lagodekhi reserve with new records for Transcaucasia and Georgia. Zootaxa 42277(1):86–98. http://www.mapress.com/j/zt/

Bača F (1994) New member of the harmful entomofauna of Yugoslavia *Diabrotica virgifera virgifera* LeConte (Coleoptera, Chrysomelidae). Zaštita Bilja 45(2): 125–131

Baselga A, Novoa F (2003) Los Chrysomelidae de los Arribes del Duero, noroeste de la Península Ibérica (Coleoptera). Nouv Rev Entomol (NS) 20(2): 117–131

Baselga A, Novoa F (2004) Coleópteros del Parque Natural de las Fragas del Eume (Galicia, noroeste de la Península Ibérica), II: Scarabaeoidea, Buprestoidea, Byrrhoidea, Elateroidea, Bostrichoidea, Lymexyloidea, Cleroidea, Cucujoidea, Tenebrionoidea, Chrysomeloidea y Curculionoidea. Boln Asoc Esp Entomol 28(1–2): 121–143

Baselga A, Novoa F (2006) Diversity of Chrysomelidae (Coleoptera) in Galicia, Northwest Spain: estimating the completeness of the regional inventory. Biodivers and Conservation 15:205–230. http://dx.doi.org/10.1007/s10531-004-6904-x

Baviera C, Biondi M (2015) The Alticini (Coleoptera: Chrysomelidae, Galerucinae) of Sicily: recent records and updated checklist. Atti della Accademia Peloritana dei Pericolanti-Classe di Scienze Fisiche, Matematiche e Naturali, 93(2), A2. http://dx.doi.org/10.1478/AAPP.932A2

Beenen R (2006) Translocation in leaf beetles (Coleoptera: Chrysomelidae). Bonn Zool Beitr 54:179–199

Beenen R (2010) Galerucinae. In Löbl, Smetana (ed) Catalogue of Palaearctic Coleoptera. Vol. 6. Apollo Books, Stenstrup, pp 443–491

Beenen R, Roques A (2010) Leaf and Seed Beetles (Coleoptera, Chrysomelidae). Chapter 8.3. in: Roques A, Kenis M, Lees D, Lopez-Vaamonde C, Rabitsch W, Rasplus JY, Roy DB (ed). Alien terrestrial arthropods of Europe. BioRisk. 4(1):267–292 http://dx.doi.org/10.3897/biorisk.4.52

Beenen R, Winkelman J, van Nunen F, Vorst O (2009) Aantekeningen over Chrysomelidae (Coleoptera) in Nederland 9. Entomol Ber 69(1):9–12

Berti N, Rapilly M (1976) Faune d’Iran—Liste d’espèces et révision du genre *Lilioceris* Reitter (Col. Chrysomelidae). Ann Soc Entomol Fr 12(1):31–73

Bieńkowski AO, Orlova-Bienkowskaja MJ (2013) New data on the composition and distribution of the genus *Leptomona* Bechyné, 1958 (Coleoptera, Chrysomelidae, Galerucinae). Entomol Review 93(7):901–903 http://dx.doi.org/10.1134/S0013873813070130

Bienkowski AO, Orlova-Bienkowskaja MJ (2015) Trophic specialization of leaf beetles (Coleoptera, Chrysomelidae) in the Volga Upland. Biol Bull 2015 42(10):863–869 http://dx.doi.org/10.1134/S1062359015100015

Bieńkowski AO (2011) Leaf beetles of European Russia. Lambert Academic Publishing, Saarbrücken

Bieńkowski AO, Orlova-Bienkowskaja MJ (2016) Key to Holarctic species of *Epitrix* flea beetles (Coleoptera: Chrysomelidae: Galerucinae: Alticini) with review of their distribution, host plants and history of invasions. Zootaxa 4175(5):401–435 http://doi.org/10.11646/zootaxa.4175.5.1

Bieńkowski AO, Orlova-Bienkowskaja MJ (2017a) World checklist of flea-beetles of the genus *Epitrix* (Coleoptera: Chrysomelidae: Galerucinae: Alticini). Zootaxa 4268(4):523–540 https://doi.org/10.11646/zootaxa.4268.4.4

Bienkowski AO, Orlova-Bienkowskaja MJ (2017b) Catalogue of locations of leaf-beetles (Chrysomelidae) of Russia and adjacent regions. Version 16.10.2017 https://www.zin.ru/Animalia/Coleoptera/rus/benkat15.htm Accessed 17 December 2017

Bieńkowski AO, Orlova-Bienkowskaja MJ (2018a) Quick spread of the invasive rose flea beetle *Luperomorpha xanthodera* (Fairmaire, 1888) in Europe and its first record from Russia (Coleoptera, Chrysomelidae, Galerucinae, Alticini). Spixiana (accepted for publication)

Bieńkowski AO, Orlova-Bienkowskaja MJ (2018b) First record of the invasive alien pest of soybeans *Medythia nigrobilineata* (Coleoptera: Chrysomelidae) in Europe. Acta Zool Bulg (accepted for publication)

Biondi B, Daccordi M, Regalin R, Zampetti M (1995) Coleoptera Polyphaga XV (Chrysomelidae, Bruchidae). In: Minelli A, Ruffo S, La Posta S (ed) Checklist delle specie della fauna italiana, 60. Calderini, Bologna

Blackburn TM, Pyšek P, Bacher S, Carlton JT, Duncan RP, Jarošík V, Wilson JRU, Richardson DM (2011) A proposed unified framework for biological invasions. Trends Ecol Evol 26(7):333–339

Bodor J (2011) Az ázsiai földibolha (*Luperomorpha xanthodera* Fairmare) megjelenése Magyarországon. Növényvédelem 47 (3): 115–116

Bolu H (2016) Southeastern Anatolia Region insect fauna I (Coleoptera I: Caraboidea; Dytiscoidea; Bostrichoidea; Chrysomeloidea; Cleroidea; Cucujoidea) of Turkey. Agriculture & Forestry 62(4):125–145 https://doi.org/10.17707/AgricultForest.62.4.16

Borowiec L, Ścibior R, Kubisz D (2011) Critical check-list of the Polish Chrysomeloidea, excluding Cerambycidae (Coleoptera: Phytophaga). Genus 22(4):579–608

Branson TF, Krysan JL (1981) Feeding and oviposition behavior and life cycle strategies of *Diabrotica*: an evolutionary view with implications for pest management. Environ Entomol 10(6):826–831

Bukejs A, Telnov D (2010) On Latvian Chrysomelinae (Coleoptera: Chrysomelidae): 4. Genus *Chrysolina* Motschulsky, 1860. Acta Zool Lituanica. 20(2):133–150. http://dx.doi.org/10.2478/v10043-010-0013-8

Butaliuk TO, Pinchuk NV, Verheles PM (2016) Analysis of distribution areas and measures to combat western corn beetle *Diabrotica virgifera* Le Conte) in the US, Europe and Ukraine. Plant Protect 4:240–249 (in Ukrainian)

Callot H (2017) Les Coléoptères du Jardin Botanique de l’Université de Strasbourg. Plus de 1000 espèces inventoriées! Bull Assoc Philomath. Alsace & Lorraine, 2014-2015(46): 111–155

Capinera JL (2001) Handbook of Vegetable Pests. Academic Press, San Diego

Chamberlin FS, Tenhet JN (1923) The tobacco flea-beetle in the southern cigar-wrapper district. U.S. Dep Agric Farmers’ Bull 1352:1–10

Champion GC (1902) VI. An Entomological Excursion to Central Spain. Ecol Entomol 50(1):115–130

Ciosi M, Miller NJ, Kim KS, Giordano R, Estoup A, Guillemaud T (2008) Invasion of Europe by the western corn rootworm, *Diabrotica virgifera virgifera*: multiple transatlantic introductions with various reductions of genetic diversity. Mol Ecol 17(16):3614–3627 https://doi.org/10.1111/j.1365-294X.2008.03866.x

Conti B, Raspi A (2007) Prima segnalazione in Italia di *Luperomorpha nigripennis* Duvivier (Coleoptera Chrysomelidae). Inf Fitopatol 57(7-8):51–52

Daccordi M, Petitpierre E (1977) Coleópteros Crisomélidos de la Sierra de Cazorla (Jaén) y descripción de una nueva especie de Clytra Laich (Col. Chrysomelidae). Misc Zool 4(1):225–236

Del Bene G, Conti B (2009) Notes on the biology and ethology of *Luperomorpha xanthodera*, a flea beetle recently introduced into Europe. Bull Insectol 62(1):61–68

Deligeorgidis PN, Ipsilandis CG, Kaltsoudas G, Sidiopoulos G, Deligeorgidis NP, Vailopoulou M, Vardiabasis A (2007) Chemical control of *Thrips tabaci, Epitrix hirtipennis* and *Myzus persicae* in tobacco fields in Northern Greece. J Entomol 4(6):463–468

Deseö KV, Balbiani A, Sannino L, Zampfelli G (1993) Zur Biologie und biologischen Bekämpfung des Tabakkäfers, *Epithrix hirtipennis* Melsh.(Col., Chrysomelidae) in Italien. Anz Schädlingskunde Pflanzenschutz Umweltschutz 66:26–29

Döberl M (1994a) Bemerkenswerte Alticinenfunde aus Westeuropa. Entomol Nachr Ber 38:179–182

Döberl M (1994b) Auff ällige Ausbreitung einiger Alticinen-Arten in Westeuropa (Coleoptera, Chrysomelidae, Alticinae). In: Verhandlungen des 14. Internationalen Symposiums für Entomofaunistiek in Mitteleuropa, SIEEC, München, pp. 276–280

Döberl M (2010) Alticinae. In Löbl, Smetana (ed) Catalogue of Palaearctic Coleoptera. Vol. 6. Apollo Books, Stenstrup, pp 491–563

Döberl M, Sprick P (2009) *Luperomorpha* Weise, 1887 in Western Europe (Coleoptera, Chrysomelidae, Alticinae). Entomol Bl Biol Systemati Käfer 105:51–56

Doguet S (2008) Présence en France de *Luperomorpha nigripennis* Duvivier, 1892 (Col. Chrysomelidae, Alticinae). Le Coléoptériste 11(1):62–63

Dolgin MM, Bieńkowski AO (2011) Leaf-beetles (Coleoptera, Chrysomelidae). Fauna of European North-East of Russia. Vol. 8, Part.3, Nauka, Saint Petersburg (in Russian)

Ekiz AN, Şen İ, Aslan EG, Gök A (2013) Checklist of leaf beetles (Coleoptera: Chrysomelidae) of Turkey, excluding Bruchinae. J Nat His 47(33-34):2213–2287. http://dx.doi.org/10.1080/00222933.2012.763069

EPPO (2017) PQR – EPPO database in quarantine pests. https://www.eppo.int/DATABASES/pqr/pqr.htm Accessed 10 December 2017

Fagot J, Libert PN (2016) Entretiens sur les Chrysomelidae de Belgique et des régions limitrophes 6. *Luperomorpha xanthodera* (Fairmaire, 1888), espèce nouvelle pour la faune belge (Chrysomelidae, Alticinae). Faunistic Entomol 69:81–82

Friedman ALL (2016) Rosemary beetle *Chrysolina americana*: A new invasive leaf beetle (Coleoptera: Chrysomelidae: Chrysomelinae) in Israel. Israel J Entomol 46:87–91

Furth DG, Savini V (1996) Checklist of the Alticinae of Central America, including Mexico (Coleoptera: Chrysomelidae). Insecta Mundi 10(1-4):45–68

García-Ocejo A, Gurrea P (1995) Los crisomélidos (Coleoptera: Chrysomelidae) de la sierra de Guadarrama (España central). Análisis biogeográfico. Boln Asoc Esp Ent 19(3-4):51–68

GBIF (2017a) Leptomona erythrocephala. Occurrence Download doi: 10.15468/dl.ykri9w Accessed via GBIF.org on 24 Oct 2017

GBIF (2017b) Chrysolina americana. Occurrence Download doi:10.15468/dl.4vzyho Accessed via GBIF.org on 26 Oct 2017

GBIF (2017c) Epitrix hirtipennis. Occurrence Download doi:10.15468/dl.0cs3vw Accessed via GBIF.org on 29th Oct 2017

GBIF (2017d) Diabrotica virgifera. Occurrence Download doi:10.15468/dl.ek5gzv Accessed via GBIF.org on 28 Oct 2017

Geiser E, Bernhard M (2012) Der Flohkäfer *Luperomorpha xanthodera* (Fairmaire, 1888) (Alticinae, Chrysomelidae) Erstnachweis für Österreich in einem Salzburger Garten. Newsl Salzburger Entomol Arbeitsgemeinschaft 3–4:1-3

Gerber E, Hinz HL, Fife D, Cristofaro M, Cristina FD, Lecce F, Paolini A, Dolgovskaya M (2012) Biological control of perennial pepperweed, *Lepidium latifolium*. Annual Report 2011 CABI Ref: VMO1732 Issued February 2012 www.cabi.org

Gerber E, Schaffner U, Gassmann A, Hinz HL, Seier M, Müller-Schärer H (2011) Prospects for biological control of *Ambrosia artemisiifolia* in Europe: learning from the past. Weed Rese 51(6):559–573

Gorodkov KB (1981) Maps 73-125. In: K.B. Gorodkov (ed) Ranges of Insects of the European Part of the USSR: An Atlas. Nauka, Leningrad (in Russian)

Gourves J, Samuelson GA (1979) Les Chrysomelidae de Tahiti (Coleopteres). Pac Insects 20(4):410–415

Gruev B, Döberl M (1997) General distribution of the flea beetles in the Palaearctic Subregion (Coleoptera, Chrysomelidae: Alticinae). Scopolia: J Slovenian Mus Nat Hist 37:1–496

Gruev B, Döberl M (2005) General distribution of the flea beetles in the Palaearctic Subregion (Coleoptera, Chrysomelidae: Alticinae). Supplement. Pensoft, Sofia-Moscow

Gruev B (2006) The leaf beetles (Coleoptera: Chrysomelidae) of the Pirin Mountain (Bulgaria). Hist Nat Bulg 17:51–79

Guskova EV (2013) Leaf beetles (Coleoptera, Chrysomelidae) of Tigirekskiy Nature Reserve (North-West Altai, Russia). Subfamilies: Chrysomelinae, Galerucinae. Alticinae Bul Altai State Agric Univ 2(100):66–72 (in Russian)

Gök A, Çilbiroglu EG, Ayvaz Y, Yildirim MZ (2002) Two new records for the Turkish flea beetle fauna: *Phyllotreta reitteri* Heik., 1911 and *Epitrix dieckmanni* Mohr, 1968 (Coleoptera, Chrysomelidae, Alticinae). Israel J Zoology 48(3):254–255

Harada H, Takizawa H (2012) Occurrence of *Epitrix hirtipennis* (Melsheimer) (Coleoptera: Chrysomelidae), an Alien Insect Pest, in Japan. Jpn J Appl Entomol Zool 56(3):117–120

Hegi G (1986) Spermatophyta, Band IV Teil 1. Angiospermae, Dicotyledones 2. In: Illustrierte Flora von Mitteleuropa. Paul Parey, Berlin, pp. 126–131

Heikertinger F (1941) Bestimmungstabelle der palaearktischen *Phyllotreta*-Arten Bestimmungstabellen europeisher Kafer. 82 Chrysomelidae 5. Subfam. Halticinae. – Gatt. *Phyllotreta* Steph. Koleopterologische Rundsch 27:15–64

Heinig U, Schöller M (2017) Rote Liste und Gesamtartenliste der Blattkäfer (Coleoptera: Chrysomelidae und Megalopodidae) von Berlin. Techische Universität Berlin http://dx.doi.org/10.14279/depositonce-5855

Hemerik L, Busstra C, Mols P (2004) Predicting the temperature-dependent natural population expansion of the western corn rootworm, *Diabrotica virgifera*. Entomol Experimentalis Applicata 111(1):59–69 http://dx.doi.org/10.1111/j.0013-8703.2004.00150.x

Hilburn DJ, Gordon RD (1989) Coleoptera of Bermuda. Florida Entomologist 72(4):673–692 http://dx.doi.org/10.2307/3495046

Hinz HL, Gerber E, Cristofaro M, Tronci C, Seier MK, Korotyaev BA, Gültekin L, Williams L, Schwarzländer M (2008) All against one: first results of a newly formed foreign exploration consortium for the biological control of perennial pepperweed. In: Proceedings of the XII international symposium on biological control of weeds. CAB International, Wallingford, pp. 154–159

Israelson G (1985) Notes on the coleopterous fauna of the Azores, with description of new species of *Athete* Thomson. Bol Mus Munic Funchal 37:5–19

Ivanchik EP, Izhevsky SS (1981) The history of Colorado potato beetle, *Leptinotarsa decemlineata* Say dispersal and its current range. In: Ushatinskaya RS (ed) The Colorado Potato Beetle, Leptinotarsa decemlineata Say, Nauka Publishers, Moscow, pp. 11–26 (in Russian)

Johnson C (1963) *Chrysolina americana* (L.) (Col. Chrysomelidae) in Britain. Entomologist’s Mon Mag 99:228–229

Johnson C, Booth RG (2004) *Luperomorpha xanthodera* (Fairmaire): a new British Flea Beetle (Chrysomelidae) on Garden Centre Roses. The Coleopterist 13(4):81–86

Jolivet P (1995) Réflexions sur les plates-hôtes des Chrysomélides (Col.). L’Entomologiste 51(2):77–93

Khlyap LA, Bobrov VV, Warshavsky AA (2010) Biological invasions on Russian territory: Mammals. Rus J Biol Invasions 1(2):127–140. http://dx.doi.org/10.1134/S2075111710020128

Kim OK, Ishikawa T, Yamada Y, Sato T, Shinohara H, Takahata K (2017) Incidence of pests and viral disease on pepino (*Solanum muricatum* Ait.) in Kanagawa Prefecture, Japan. Biodiversity data journal, (5). Biodivers Data J. (5): e14879. Published online 2017 Aug 22. http://dx.doi.org/10.3897/BDJ.5.e14879

Kippenberg H (2010) Chrysomelinae. In Löbl, Smetana (ed) Catalogue of Palaearctic Coleoptera. Vol. 6. Apollo Books, Stenstrup, pp 390–443

Kondratenko NN (2012) Production of soybean in Krasnodar Krai. http://7law.info/krasnodar/act2d/u679.htm. Accessed 17 December 2017 (in Russian)

Kovalev OV (2002) Formaton of solitary population waves in invasions of organisms and evolution of biosphere. Evolutionary Botany, Vol. 2. Proceedings of the 2nd international conference Problem of species and speciation (Tomsk, 24-26 October 2001). pp 65–81 (in Russian)

Kovalev OV, Tyutyunov YuV, Arkhipova OE, Kachalina NA, Iljina LP, Titova LI (2015) On assessment of the large-scale effect of introduction of the ragweed leaf beetle *Zygogramma suturalis* F. (Coleoptera, Chrysomelidae) on the phytocenoses of South Russia. Entomol Rev 95(1):1–14. http://dx.doi.org/10.1134/S0013873815010017

Kovalev OV, Medvedev LN (1983) Theoretical Foundations of Introduction of Ragweed Leaf Beetles of the Genus Zygogramma Chevr. (Coleoptera, Chrysomelidae) for the Purpose of Biological Control of Ragweeds in the USSR. Entomologicheskoe Obozrenie 62(1):17–32 (in Russian)

Kovalev OV, Tyutyunov YV, Iljina LP, Berdnikov SV (2013) On the efficiency of introduction of American insects feeding on the common ragweed (*Ambrosia artemisiifolia* L.) in the south of Russia. Entomol Rev 93(8):962–973 http://dx.doi.org/10.1134/S0013873815010017

Koyama T (1940) On the Hibernation of *Paraluperodes suturalis nigrobilineatus* Motsch. Kontyu 2(6):256–259

Kozłowski MW, Legutowska H (2014) The invasive flea beetle *Luperomorpha xanthodera* (Coleoptera: Chrysomelidae: Alticinae), potentially noxious to ornamental plants – first record in Poland. J Plant Prot Res 54(1):106–107 http://dx.doi.org/10.2478/jppr-2014-0017

Krsteska V, Dimeska V, Stojanoski P (2009) *Epitrix hirtipennis* Melsh on tobacco. In: Abstracts of presentations made at the 2009 Coresta Joint Meeting of the Agronomy and Phytopatology Study Groups, Agro/Phyto, Rovinj, p. 15

Krsteska V, Spirkoski A (2017) A contribution to quantitative representation and distribution of *Epitrix hirtipennis* (Melsheimer, 1847) (Coleoptera: Chrysomelidae, Alticini) on tobacco. Acta Entomol Serbica, 22. http://dx.doi.org/10.5281/zenodo.857850

Lombaert E, Ciosi M, Miller NJ, Sappington TW, Blin A, Guillemaud T (2017) Colonization history of the western corn rootworm (*Diabrotica virgifera virgifera*) in North America: insights from random forest ABC using microsatellite data. bioRxiv, 117424 http://dx.doi.org/10.1101/117424

Lopatin IK (2010) Leaf beetles (Insecta, Coleoptera, Chrysomelidae) of Central Asia. BSU, Minsk (in Russian)

Lopatin IK (1977) Leaf beetles of Middle Asia and Kazakhstan: A key. Leningrad (in Russian) Lykouressis DP (1991) *Epitrix hirtipennis*, a new pest of tabacco in Greece, with notes of its morphology, bioecology and control. Entomologica Hellenica 9:81–85

MacLeod A (2002) CSL pest risk analysis for *Chrysolina americana*. 5 pp. https://secure.fera.defra.gov.uk/phiw/riskRegister/viewPestRisks.cfm?cslref=8564 Accessed 17 December 2017

Maican S (2005) Checklist of Chrysomelidae (Coleoptera) of Romania. Travaux Mus Natl Hist Nat Grigore Antipa 48:119–136

Majka CG, LeSage L (2008) Introduced leaf beetles of the Maritime provinces, 5: the lily leaf beetle, *Liloceris lilii* (Scopoli) (Coleoptera, Chrysomelidae). Proc Soc Entomol Wash 110(1):186–195

Manole T, Chireceanu C, Teodoru A (2017) The broadening of distribution of the invasive species *Diabrotica virgifera virgifera* Leconte in the area of Mubtenia region under specific climatic and trophic conditions. Scientific Papers. Series A. Agronomy 60:495–499

Martin WD, Herzog GA (1987) Life history studies of the tobacco flea beetle, *Epitrix hirtipennis* (Melsheimer) (Coleoptera: Chrysomelidae). J Entomol Sci:237–244

Maslyakov VYu, Izhevsky SS (2011) Alien phytophagous insect invasions in the European part of Russia. IGRAS, Moscow (in Russian)

Matsishina NV (2011) On the biology of Colorado potato beetle *Leptinotarsa decemlineata* Say, 1824 (Coleoptera: Chrysomelidae) in the south of the Russian Far East. Euroasian Entomol J 10(3):330–336 (in Russian)

Medvedev LN (1992) Chrysomelidae – Leaf-beetles. In: Key to insects of the Far East of USSR. 3(2), Nauka, St.-Petersburg, pp. 533–602 (in Russian)

Medvedev LN, Roginskaya EYa (1988) Catalogue of host plants of leaf beetles of the USSR. Moscow, A.N. Severtsov Institute of Evolutionary Morphology and Ecology, Moscow (in Russian)

Moseiko AG (2010) Estimation of agricultural significance of Chrysomelidae (Coleoptera) species damaging soya in the Far East. Plant Protect News 1:42–47 (in Russian)

Mosyakin SA (1987) Ecological-faunistic survey of leaf beetles of Crimea. In: Abstracts of the 3rd congress of Ukrainian Entomol Sci, Kiev, pp 129–130 (in Russian)

Nishida GM ed. (2002) Hawaiian Terrestrial Arthropod Checklist. 4th edn. Bishop Museum Technical Report 22, Hawaii Biological Survey, Bishop Museum, Honolulu

Ntibiyoboka J (2014) Economics of smallholder tobacco production and marketing in Mpanda District. Doctoral dissertation, Sokoine University of Agriculture

Ogilvie L (1924) Preliminary report of the plant pathologist for the period September 27th to December 31st, 1923. Bermuda Department of Agriculture Annual Report, 1923:28–34.

Ogloblin DA (1936) Insects. Coleoptera. Chrysomelidae, Galerucinae. Fauna of USSR 26(1) Academia of Sciences of USSR Publ., Moscow and Leningrad. (in Russian)

Orlova-Bienkowskaja MJ (2013a) Disjunctive area of *Chrysolina eurina* (Frivaldszky, 1883) (Coleoptera: Chrysomelidae: Chrysomelinae). Caucasian Entomol Bul 9(1):102–107

Orlova-Bienkowskaja MJ (2013b) Dynamics of the range of lily leaf beetle (*Lilioceris lilii*, Chrysomelidae, Coleoptera) indicates its invasion from Asia to Europe in the 16th-17th century. Rus J Biol Invasions 4(2):93–104. http://dx.doi.org/10.1134/S2075111713020082

Orlova-Bienkowskaja MJ (2014) First record of the tobacco flea beetle *Epitrix hirtipennis* Melsheimer [Coleoptera: Chrysomelidae: Alticinae] in Russia. EPPO Bulletin 44(1):44–46. http://dx.doi.org/10.1111/epp.12092

Orlova-Bienkowskaja MJ (2016) Is it possible to distinguish alien species of beetles (Coleoptera) from native ones? Entomol Rev 96(3):318–331. http://dx.doi.org/10.1134/S001387381603009X

Orlova-Bienkowskaja MJ (2017) Main trends of invasion processes in beetles (Coleoptera) of European Russia. Rus J Biol Invasions 8(2): 143–157. http://dx.doi.org/10.1134/S2075111717020060

Özdikmen H, Mercan N, Cihan N, Kaya G, Topcu NN, Kavak M (2014) The importance of superfamily Chrysomeloidea for Turkish biodiversity (Coleoptera). Munis Ent Zool 9:17–45

Özdikmen H, Şahin DC, Bal N (2017) New food plants and new records of two species of *Epitrix* Foudras in Turkey (Chrysomelidae: Galerucinae: Alticini). Ent Zool 12(1):309–312

Pasqual C, Kovar I, Beenen R, van Vondel BJ, Audisio P (2017) Fauna Europaea: *Chrysolina (Taeniochrysea) americana*. Fauna Europaea version 2017.12. https://fauna-eu.org/cdmdataportal/taxon/2f547fb0-ef3a-44a7-b884-4f868b3847ff Accessed 17 December 2017

Petitpierre E (1981) Chrysomelidae (Col.) de la sierra de Albarracín (Teruel). Boln Asoc Esp Entomol 4:7–18

Petitpierre E (1997) Los Chrysomelidae (Coleoptera) del Moncayo (Aragón). Zapateri, Rev Aragonesa Entomol 7:273–280

Petitpierre E (2005) Listado de Chrysomelidae (Coleoptera) de Asturias y Cantabria. Boln Asoc Esp Ent 29:51–72

Petitpierre E (2007) Catàleg dels coleòpters crisomèlids de Catalunya V. Hispinae i Cassidinae, i llista actualitzada de totes les espècies de la família. Butl Inst Catalana Hist Nat 75:61–83

Petitpierre E, Bastazo G, Vela JM (2011) Estudio faunístico de los crisomélidos de la provincia de Cádiz, España (Coleoptera, Chrysomelidae). Zool Baetica 22:137–170

Petitpierre E, Daccordi M (2013) Chrysomelidae (Coleoptera) de la sierras del Altiplano de Granada (Granada, Andalucía). Zool Baetica 24:53–78

Petitpierre E, Sacarés A, Jurado-Rivera JA (2017) Updated checklist of Balearic leaf beetles (Coleoptera: Chrysomelidae). Zootaxa 4272(2):151–177 https://doi.org/10.11646/zootaxa.4272.2.1

Plotnikova TV (2014) New pest in tobacco agrocenosis of Russia. Plant Protect Quar J 5:41–42 (in Russian)

Popova EN (2014) The influence of climatic changes on range expansion and phenology of the Colorado potato beetle (*Leptinotarsa decemlineata*, Coleoptera, Chrysomelidae) in the territory of Russia. Entomol Rev 94(5):643–653. https://doi.org/10.1134/S0013873814050017

Pyšek P, Sádlo J, Mandák B (2002) Catalogue of alien plants of the Czech Republic. Preslia 74:97–186

Pyšek P, Meyerson LA, Simberloff D (2017) Introducing “Alien Floras and Faunas”, a new series in Biological Invasions. Biol Invasions, Published online. https://doi.org/10.1007/s10530-017-1648-1

Raspudić E, Brmež M, Budimir A, Pleša Z, Zdeličan J (2015) *Epitrix hirtipennis* (Melsheimer, 1847.) duhanov buhač, novi član entomofaune hrvatske. Book of abstracts 12th Croatian Biological Congress with International Participation, Klobučar, G., Kopjar, N., Gligora Udovič, M., Lukša, Ž., Jelić, D. (ur.), Hrvatsko biološko društvo, Zagreb, 97–97 http://bib.irb.hr/prikazi-rad?rad=782348

Reitter E (1886) Coleopteren aus Europa und den angrenzenden Ländern mit Bemerkungen über bekannte Arten. Zweiter Thile. Dtsch Entomol Z 30:67–72

Reznik SYa, Spasskaya IA (2005) Population densities of the ragweed leaf beetle *Zygogramma suturalis* F. (Coleoptera: Chrysomelidae) in the North Caucasus in 2005. Proc Rus Entomol Soc St. Petersburg 77:267–271 (in Russian)

Reznik SYa, Spasskaya IA, Dolgovskaya MY, Volkovitsh MG, Zaitzev VF (2008) The ragweed leaf beetle *Zygogramma suturalis* F. (Coleoptera: Chrysomelidae) in Russia: current distribution, abundance, and implication for biological control of common ragweed, *Ambrosia artemisiifolia* L. In: Julien MH et al. (ed) Proc. of the XII Intern. Symp. On Biological Control of Weeds CABI, Wallingford, pp. 614–619

Richardson DM, Pyšek P, Rejmanek M, Barbour MG, Panetta FD, West CJ (2000) Naturalization and invasion of alien plants: concepts and definitions. Diversity and Distributions 6(2):93–107

Riley EG, Clark SM, Seeno TN (2003) Catalog of the leaf beetles of America North of Mexico (Coleoptera: Megalopididae, Orsodacnidae and Chrysomelidae, excluding Bruchinae). Coleopterists Society, Sacramento

Royal horticultural society (2014) Rosemary beetle. https://www.rhs.org.uk/advice/profile?pid=555 Accessed 17 December 2017

Samuelson GA (1973) Alticinae of Oceania (Coleoptera, Chrysomelidae). Pacific Insects Monograph. Vol. 30 Bishop Museum, Honolulu

Sannino L, Balbiani A (1990) Attuali possibilità di controllo di *Epitrix hirtipennis* in Italia. L’Informatore Agrario 46 Suppl 13:17–20

Sannino L, Balbiani A, Espinosa B (1984) Un nuovo fitofago devasta il tabacco nel beneventano: *Epithrix hirtipennis* Melsh. (Coleoptera: Chrysomelidae). L’Informatore Agrario 29:55–57

Sannino L, Balbiani A, Biondi M (1985) *Epithrix hirtipennis* (Melsheimer, 1847): Considerazioni tassonomiche, ecologiche ed etologiche. In: Atti XIV Congresso Nazionale Italiano di Entomologia, Palermo, Erice, Bagheria, pp. 285–292

Schmidt MH, Lefebvre G, Poulin B, Tscharntke T (2005) Reed cutting affects arthropod communities, potentially reducing food for passerine birds. Biol Conserv 121:157–166 http://dx.doi.org/10.1016/j.biocon.2004.03.032

Sergeev MYe (2008) Biology and perspectives of use of *Zygogramma suturalis* (F.) (Coleoptera, Chrysomelidae) in the south-east of the Ukraine for control of *Ambrosia artemisiifolia*. In: Proceedings of the 3rd international scientific conference «Restoration of disturbed natural ecosystems » (Donetsk, 7-9 October 2008). Tsifrovaya tipografia, Donetsk, pp. 496–501 (in Russian)

Sergeev MYe (2010) Leaf-beetles (Coleptera: Chrysomelidae) of sand-coquinal terrace of Azov sea, Ukraine. Caucasian Entomol Bull 6(2):161–170 (in Russian)

Sergeev MYe (2011) Contribution to the fauna of leaf-beetles (Coleoptera, Chrysomelidae) of Ukrainian Steppe Nature Reserve, with a review of materials from other regions of Ukraine. Ukrainska Entomofaunistyka 2(4): 1–29 (in Russian)

Sergeev MYe (2012) Usage of *Zygogramma suturalis* F. (Coleoptera, Chrysomelidae) against *Ambrosia artemisiifolia* L. in the South-East of Ukraine. In: Industrial Botany Collection of Scientific works, 12. Donetsk Botanical Garden of National Academy of Sciences of Ukraine, Donetsk, pp 49–52 (in Russian)

Sergeev MYe (2013) The First Find of *Zygogramma suturalis* (Coleoptera, Chrysomelidae, Chrysomelinae) in the Right-Bank Ukraine. Ukrainska Entomofaunistyka 4(1):49–51 (in Russian)

Sharp D (1900) Coleoptera Phytophaga. In: Fauna Hawaiiensis. Vol. 2. The University press, Cambridge, pp. 91–116

Spasov D, Spasova D, Atanasova B, Serafimova M (2013) Pest insects at tobacco (*Nicotiana tabacum* L.) in Strumica region, Republic of Macedonia. Sci Works 2(1): 193–196

Staines CL, Whittington AE (2003) Chrysomelidae (Coleoptera) types in the Royal Museum of Scotland Collection. Zootaxa 192(1):1–8

Takei M, Nakamura M, Hamada Y, Ikeda A, Mitsuya S, Suralta R, Yamauchu A (2014) Assessment of damage caused by two-striped leaf beetle (*Medythia nigrobilineata* Motschulsky) larval feeding of root nodules in soybean and its control during furrow cultivation at seeding time. Plant Prod Sci 17(3):276–283 http://dx.doi.org/10.1626/pps.17.276

Toepfer S, Li H, Pak SG, Son KM, Ryang YS, Kang SI, Han R, Holmes K (2014) Soil insect pests of cold temperate zones of East Asia, including DPR Korea: A review. J Pest Sci 87:567–595 http://dx.doi.org/10.1007/s10340-013-0540-8

Trenchev G, Tomov R (2000) Tobacco flea beetle *Epitrix hirtipennis* (Melsheimer) (Coleoptera, Chrysomelidae), a new serious pest on tobacco in Bulgaria. Yearbook for Plant Protection, Skopje 11:61–64

Trepashko LI, Nadtocheva SV (2013) Invasion of the western corn rootworm to the territory of Belarus. Plant Protection Quar J 5:40–41 (in Russian)

Vincent R, Doguet S (2011) L’altise *Luperomorpha xanthodera* (Fairmaire, 1888) poursuit son expansion en France (Coleoptera Chrysomelidae). Bull Mens Soc Linnéenne Lyon 80(9-10):218–220

VINNKR (2012) Western corn rootworm is already in Russia. http://agropost.ru/rastenievodstvo/bolezni-vrediteli/zapadniy-kukuruzniy-zhuk-uzhe-v-rossii.html Accessed 17 December 2017 (in Russian)

Vińolas A, Batet JM, Soler J (2016) Noves o interessants localitzacions d’espècies de coleòpters per a la península Ibèrica i les illes Canàries (Coleoptera). Butll Inst Catalana d’Història Nat 80:101–112

von Heyden L, Allard E (1870) Entomologische reise nach dem südlichen Spanien: der Sierra Guadarrama und Sierra Morena. Portugal und den Cantabrischen gebirgen. https://books.google.ru/books?hl=ru&lr=&id=FXBTAAAAcAAJ&oi=fnd&pg=PA1&dq=Monolepta+erythrocephala&ots=xTHUCtIdA8&sig=C1qGmp2Z_XhS33pJkRdQP1a6BGY&redir_esc=y&#x0023;v=onepage&q=Monolepta%20erythrocephala&f=false Accessed 17 December 2017

Warchalowski A (2010) The Palearctic Chrysomelidae. Identificacion keys. Vol. 1, 2. Natura optima dux Foundation, Warszawa

Waterhouse DF (1997) The major invertebrate pests and weeds of agriculture and plantation forestry in the southern and western Pacific. ACIAR Monograph No. 44. ACIAR, Canberra

Webster E, Cameron RWF, Culham A (2017) Gardening in a changing climate. Royal Horticultural Society, UK

Yaroshenko VA (1994) Leaf-beetles in natural and anthropogenic ecosystems of the Northern Caucasus. Dissertation, Moscow State University (in Russian)

Zhanglin C, Junyi G, Dongfeng J, Zhenjing R (1997) A Study on leaf-feeding insect species on soybeans in Nanjing area. Soybean Sci 16:12–20

